# Brain-Restricted mTOR Inhibition with Binary Pharmacology

**DOI:** 10.1101/2020.10.12.336677

**Authors:** Ziyang Zhang, Qiwen Fan, Xujun Luo, Kevin J. Lou, William A. Weiss, Kevan M. Shokat

## Abstract

On-target-off-tissue drug engagement is an important source of adverse effects that constrains the therapeutic window of drug candidates. In diseases of the central nervous system, drugs with brain-restricted pharmacology are highly desirable. Here we report a strategy to achieve inhibition of mTOR while sparing mTOR activity elsewhere through the use of a brain-permeable mTOR inhibitor RapaLink-1 and brain-impermeable FKBP12 ligand RapaBlock. We show that this drug combination mitigates the systemic effects of mTOR inhibitors but retains the efficacy of RapaLink-1 in glioblastoma xenografts. We further present a general method to design cell-permeable, FKBP12-dependent kinase inhibitors from known drug scaffolds. These inhibitors are sensitive to deactivation by RapaBlock enabling the brain-restricted inhibition of their respective kinase targets.

---

Administration of a small molecule drug often leads to systemic pharmacological effects that contribute to both efficacy and toxicity and define its therapeutic index. While off-target effects may be mitigated by chemical modifications that improve the specificity of the drug, on-target-off-tissue toxicity represents a unique challenge that requires precise control over tissue partitioning. On-target-off-tissue toxicities affect commonly used drugs such as statins (myopathy caused by HMG CoA reductase inhibition in skeletal muscle)^1^ and first-generation antihistamines (drowsiness caused by H_1_ receptor blockade in the brain)^2^ and can sometimes preclude the safe usage of otherwise effective therapeutic agents. Chemical inhibitors of the mechanistic target of rapamycin (mTOR) provide a case in point. Although mTOR inhibitors have shown efficacy in treating a number of central nervous system (CNS) diseases including tuberous sclerosis complex (TSC)^3,4^, glioblastoma (GBM)^5,6^, and alcohol use disorder (AUD)^7–9^, systemic mTOR inhibition is associated with a variety of dose-limiting adverse effects: immune suppression, metabolic disorders, and growth inhibition in children^10,11^. If the pharmacological effects of these mTOR inhibitors could be confined to the CNS, their therapeutic window could be substantially widened. Here we present a chemical strategy that allows brain-specific mTOR inhibition through the combination of two pharmacological agents – a brain-permeable mTOR inhibitor (RapaLink-1) whose function requires the intracellular protein FK506-binding protein 12 (FKBP12) and a brain-impermeant ligand of FKBP12 (RapaBlock). When used in a glioblastoma xenograft model, this drug combination drove tumor regression without detectable systemic toxicity. We further demonstrate that this strategy can be adapted to achieve brain-specific inhibition of other kinase targets by developing a method to convert known kinase inhibitors into FKBP12-dependent formats.

Therapeutic targeting of mTOR kinase can be achieved both allosterically and orthosterically. First-generation mTOR inhibitors rapamycin and its analogs (rapalogs) bind to the FK506 rapamycin binding (FRB) domain of mTOR as a complex with the intracellular protein FKBP12, resulting in substrate-dependent allosteric inhibition of mTOR complex 1^12–14^. Second-generation mTOR kinase inhibitors (TORKi) directly bind in the ATP pocket of mTOR and inhibit the activity of both mTOR complex 1 and complex 2^15–17^. The third generation mTOR inhibitor RapaLink-1 is a bitopic ligand that simultaneously engages the ATP pocket and an allosteric pocket (the FRB domain) of mTOR, achieving potent and durable inhibition of its kinase activity^18^. Despite its large molecular weight (1784 Da), RapaLink-1 is cell- and brain-permeant and has shown enhanced *in vivo* efficacy in driving glioblastoma regression compared to earlier mTOR inhibitors^6^.

Because RapaLink-1 contains a TOR kinase inhibitor (TORKi) moiety (Fig. 1a, grey shades) that may directly bind in the ATP pocket, we wondered whether the interaction of RapaLink-1 with FKBP12 is essential for its inhibition of mTOR. We performed *in vitro* kinase assay with purified mTOR protein and found that RapaLink-1 exhibited identical IC50 values as MLN0128 (the TORKi portion of RapaLink-1) and that inclusion of 10 µM FKBP12 had no effect on its activity (Fig. 1b). Surprisingly, when we tested RapaLink-1 in cells, we observed a strong dependency on FKBP12 for mTOR inhibition. In (K562/dCas9-KRAB cells), CRISPRi-mediated knockdown of FKBP12 with two distinct single guide RNAs (sgRNAs) drastically impeded its cellular activity (Fig. 1c). This effect was even more pronounced when we used FK506 (a high affinity natural ligand of FKBP12) to pharmacologically block the ligand binding site of FKBP12. Similar FKBP12-dependence of RapaLink-1 was observed when we monitored mTOR signaling by Western Blot (Fig. 1d, see also ref^18^) – whereas 10 nM RapaLink-1 reduced P-S6 (S240/244) and P-4EBP(T37/46) to almost undetectable levels, combination of 10 nM RapaLink-1 and 10 µM FK506 showed no effect on either marker.

**Figure 1.**
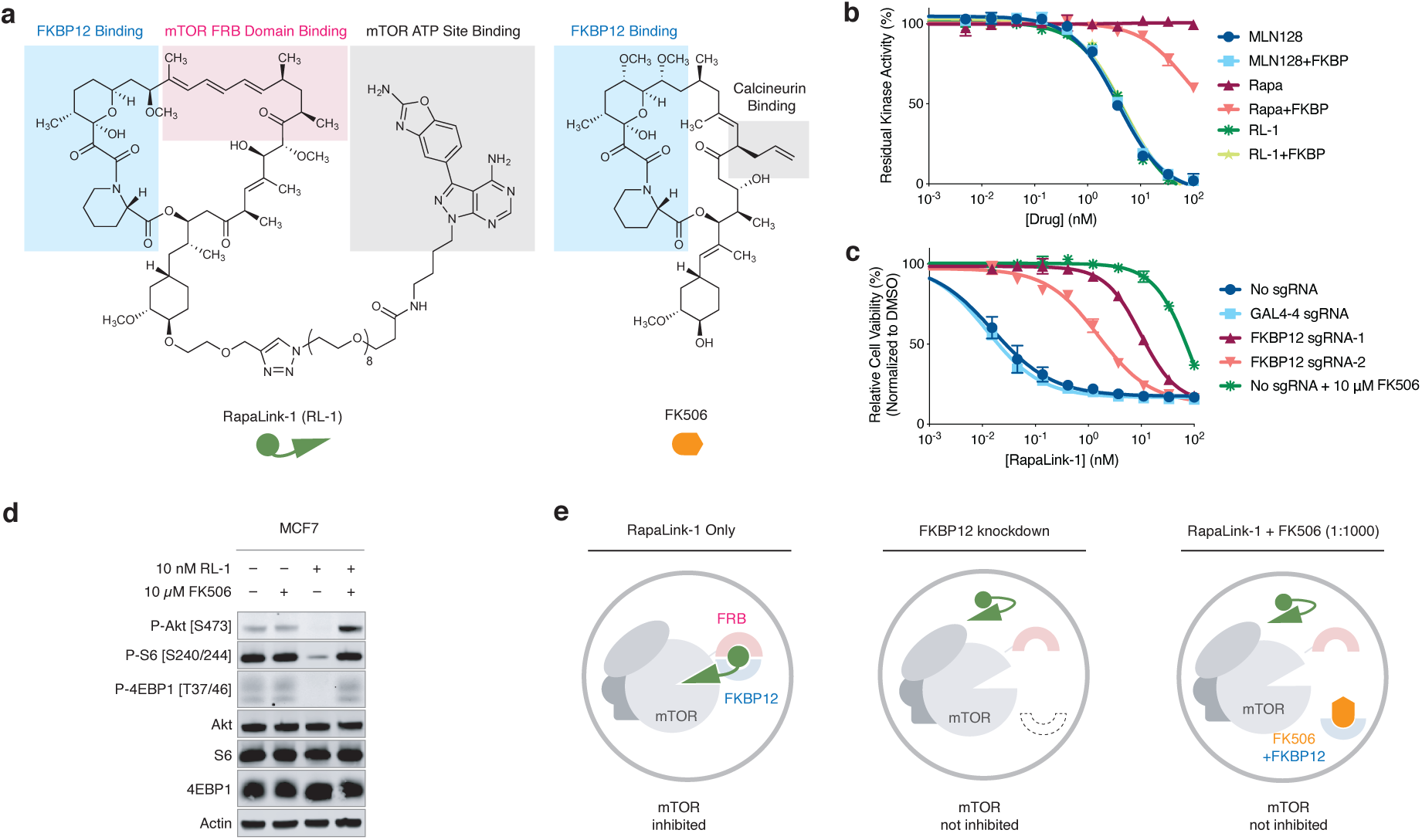
RapaLink-1 is a potent mTOR inhibitor that requires FKBP12 for its cellular activity. **a**, Chemical structures of RapaLink-1 and FK506. **b**, Inhibition of mTOR activity by MLN128, RapaLink-1 (RL-1) or Rapamycin in the presence or absence of 10 µM FKBP12 *in vitro* kinase assay. **c**, K562-dCas9-KRAB cells transduced with GAL4-4 (control) or FKBP12-targeting sgRNAs were treated with RapaLink-1 and cell proliferation was assessed after 72 h. In the last listed condition, cells were transduced with sgRNA but treated with RapaLink-1 in the presence of 10 µM FK506. **d**, Immunoblot analysis of mTOR signaling in MCF7 cells treated with DMSO, RapaLink-1, FK506, or a combination of RapaLink-1 and FK506. **e**, Schematics of the proposed working model. Grey circles indicate cellular membranes.

These results highlight that while FKBP12 is not essential for RapaLink-1 to bind and inhibit the active site of mTOR in vitro, it is required for cellular activity perhaps serving as an intracellular sink for RapaLink-1 to accumulate in the cell. We probed this possibility using a structural analog of RapaLink-1, where the TORKi moiety had been replaced with tetramethylrhodamine (RapaTAMRA, Extended Data Fig. 1). This fluorescent analog of RapaLink-1 allowed us to quantify intracellular compound concentration by flow cytometry. Consistent with previous findings with RapaLink-1^18^ and other FKBP-binding compounds^19,20^, RapaTAMRA showed high cellular retention (10-fold increase in median fluorescence intensity) even after extensive washout, but this effect was diminished by FKBP12 knockdown (Extended Data Fig. 1c). While other factors may contribute, we believe that FKBP12-mediated cellular partitioning is partly responsible for the exceptional potency of RapaLink-1.

We next considered whether the dependence of RapaLink-1 on FKBP12 could be harnessed to achieve CNS-restricted mTOR inhibition. Specifically, selective blockade of FKBP12 in peripheral tissues with a potent, brain-impermeant small molecule ligand would allow RapaLink to accumulate in the brain but not peripheral tissues, resulting in brain-specific inhibition (Fig. 2a). To identify a candidate FKBP12 ligand with the suitable permeability profile (i.e. cell-permeable but BBB-impermeable), we considered known natural and synthetic high-affinity FKBP12 ligands (e.g., FK506^21,22^, rapamycin^23,24^, SLF/Shield-1^25,26^), but all these compounds readily cross the blood-brain barrier (BBB). Therefore, we synthesized a panel of derivatives of SLF and FK506 where polar substituents (particularly hydrogen bond donors and acceptors) had been attached to the solvent exposed parts of these two molecules, a design principle directly against the empirical rules for designing BBB-permeable drugs^27–30^ (Extended Data Fig. 2). In addition, modification of the C21-allyl of FK506 has the added advantage that it abolishes the binding to calcineurin^31^, the natural target of FK506, whose inhibition causes suppression of Nuclear factor of activated T-cells (NFAT) signaling and hence immunosuppression (Fig. 2b). The majority of Shield-1 and FK506 derivatives we synthesized maintained potent FKBP12 binding (Extended Data Fig. 2), as measured by a competition fluorescence polarization assay^32^. We then screened these compounds (10 µM) in a cell-based assay, evaluating whether they can protect mTOR from inhibition by RapaLink-1 (10 nM) by monitoring phospho-S6 level by Western Blot. While none of the SLF derivatives were found to be effective, most FK506 analogs attenuated the potency of RapaLink-1 (rescuing phospho-S6 levels), with some of them completely blocking RapaLink-1 from inhibiting mTOR (Extended Data Fig. 2). Four of the most effective RapaLink-1 blocking compounds were subjected to an additional *in vivo* screen, where mTOR signaling was separately analyzed in the brain and skeletal muscle tissues of mice treated with a combination of RapaLink-1 and the candidate compound (1 mg/kg and 40 mg/kg, respectively). A pyridine *N*-oxide derivative of FK506 protected peripheral tissues from RapaLink-1 but allowed potent inhibition of mTOR in the brain (Extended Data Fig. 3). We therefore chose this compound for our further study and refer to it as “RapaBlock”.

**Figure 2.**
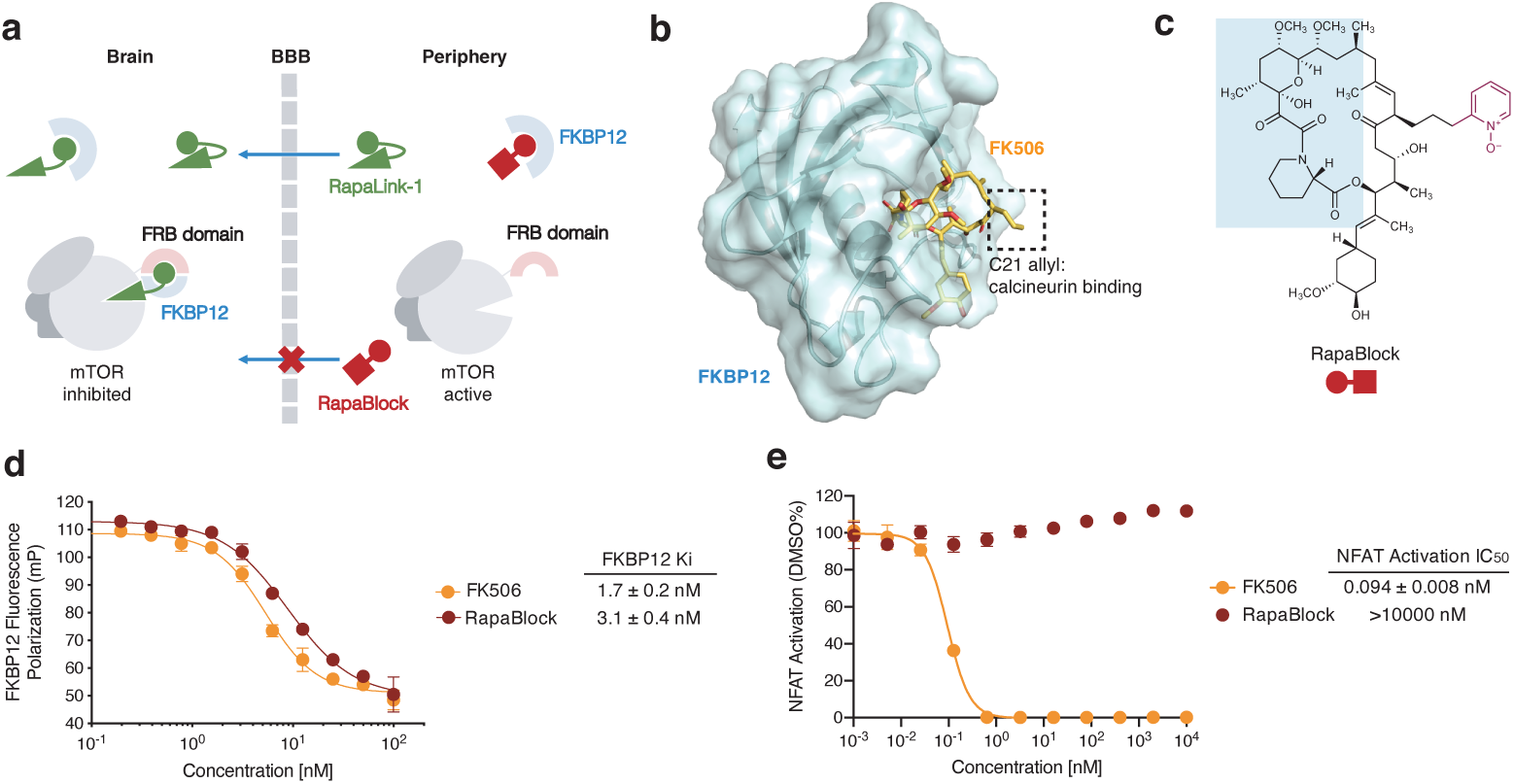
RapaBlock is a potent, non-immunosuppressive FKBP12 ligand. **a**, a proposed model to achieve brain-specific mTOR inhibition through the combination of an FKBP12-dependent mTOR inhibitor (RapaLink-1) and a brain impermeable FKBP12 ligand (RapaBlock). **b**, a FK506–FKBP12 co-crystal structure (PDB: 1FKJ) showing that the C21 allyl group is solvent exposed and its modification may lead to abolished calcineurin binding. **c**, chemical structures RapaBlock. Blue shaded areas indicate the FKBP12-binding moiety. **d**, competition fluorescence polarization assay using fluorescein-labeled rapamycin as the tracer compound (*n* = 3). **e**, Jurkat cells expressing a luciferase under the control of NFAT transcription response element were stimulated with phorbol myristate acetate and ionomycin in the presence of various concentrations of compounds (*n* = 3).

RapaBlock and FK506 bind to FKBP12 with comparable affinity (Fig. 2d, *K*_i_’s: 3.1 nM and 1.7 nM, respectively), but unlike FK506, RapaBlock does not exhibit any inhibitory activity for calcineurin (Fig. 2e). In cultured cells, RapaBlock does not affect mTOR signaling by itself (up to 10 µM) but attenuates the pharmacological effects of RapaLink-1 in a dose-dependent fashion (Fig. 3a). Interestingly, RapaBlock appears more effective at blocking rapamycin, restoring phospho-S6 signal to the same level as untreated cells at 100:1 stoichiometry (Fig. 3d). Because rapamycin and rapalogs exert immunosuppressive effects by inhibiting mTOR activity and thus suppressing cell proliferation in response to costimulatory and cytokine-mediated signals^33,34^, we asked whether RapaBlock can prevent RapaLink-1-mediated immunosuppression. We stimulated human peripheral mononuclear blood cells (PBMC) with anti-CD3 and anti-CD28 antibodies in the presence of different concentrations of RapaLink-1 and RapaBlock and assessed cell proliferation after 5 days. Whereas RapaLink-1 potently inhibited PBMC proliferation at nanomolar concentrations, addition of RapaBlock abolished this effect, shifting the IC_50_ by more than 100-fold (Fig. 3b). Though mTOR inhibition does not directly control cytokine production, we observed higher cumulative IL-2 release in cells co-treated with RapaLink-1 and RapaBlock compared to cells treated with RapaLink-1 alone, presumably as a result of greater cell proliferation (Fig. 3c). Consistent with our observation of mTOR signaling, RapaBlock appeared more effective at protecting PBMC from rapamycin-mediated proliferation inhibition (Figs. 3e, 3f). Together, these data demonstrate that by competitively binding to intracellular FKBP12, RapaBlock renders RapaLink-1 and rapamycin incapable of inhibiting mTOR, and in so doing diminishes the immunosuppressive effects of these two drugs.

**Figure 3.**
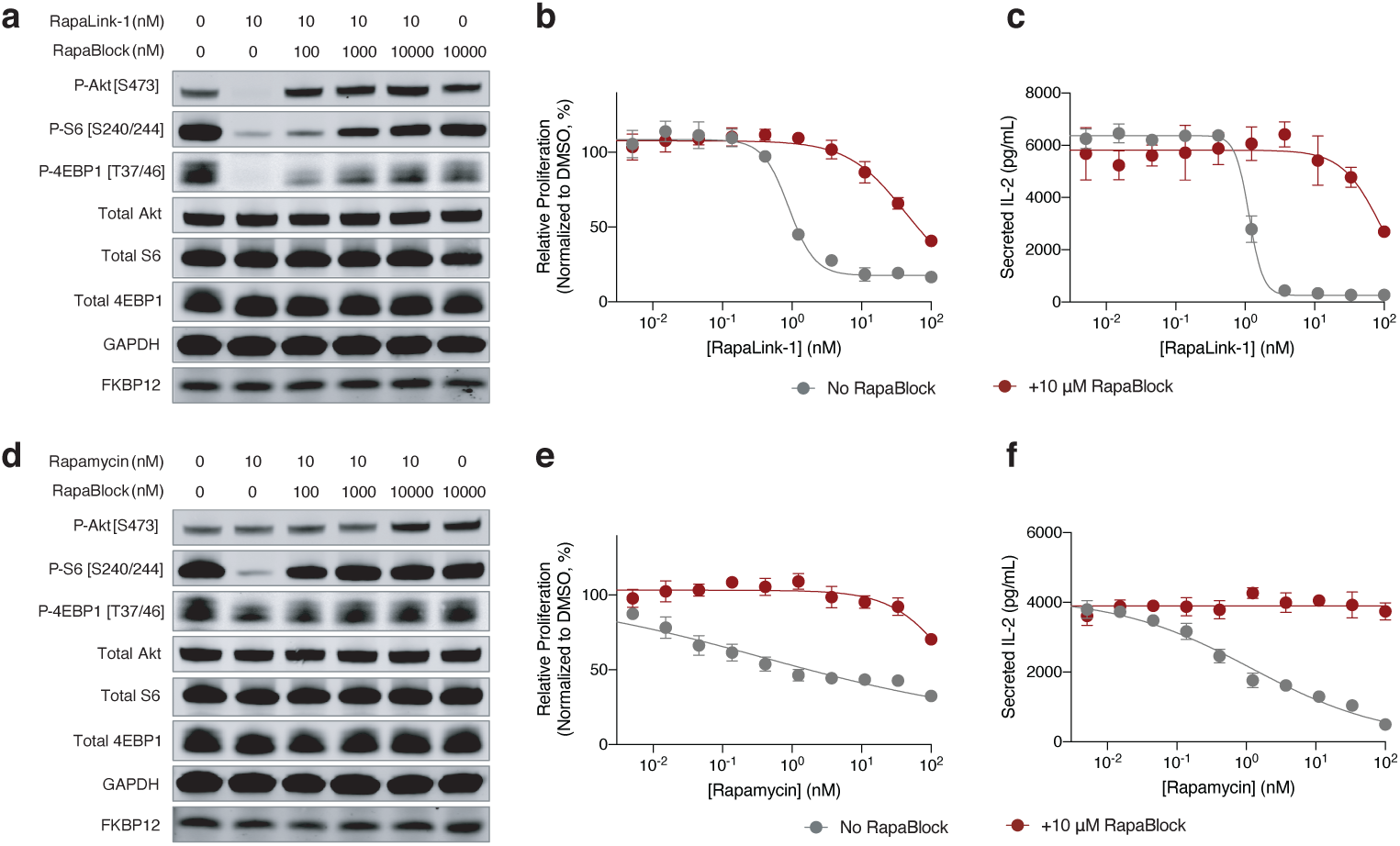
RapaBlock protects cells from mTOR inhibition by RapaLink-1 and Rapamycin. **a, d**, MCF7 cells were treated with a combination of RapaLink-1 and RapaBlock (**a**), or Rapamycin and RapaBlock (**d**) for 4 h, then phosphorylation of mTOR substrates were analyzed by immunoblotting. Results shown are representative of three independent experiments. **b, e**, Human PBMCs were stimulated with anti-CD3 and anti-CD28 in the presence of varying amounts of RapaLink-1 and RapaBlock (**b**) or rapamycin and RapaBlock (**e**), and cell proliferation was measured after 120 h. **c, f**, Interleukin-2 secretion in the culture supernatant of PBMCs in **c** and **f** was quantified by sandwich ELISA.

To examine whether the combination of RapaLink-1 and RapaBlock allows brain-specific inhibition of mTOR *in vivo*, we treated healthy BALB/c^nu/nu^ mice with RapaLink-1 (1 mg/kg) or a combination of RapaLink-1 (1 mg/kg) and RapaBlock (40 mg/kg), stimulated mTOR activity with insulin (250 mU) after 4 h or 24 h, and analyzed dissected tissues by immunoblot (Fig. 4a). RapaLink-1, when used as a single agent, potently inhibited mTOR signaling in both skeletal muscle and brain tissues, as revealed by the reduced phosphorylation of S6 and 4EBP1. The combination of RapaLink-1 and RapaBlock, however, exhibited remarkable tissue-specific effects: while mTOR activity in the brain was inhibited at a comparable level to mice treated with RapaLink-1 only, mTOR activity in skeletal muscle was not affected.

**Figure 4.**
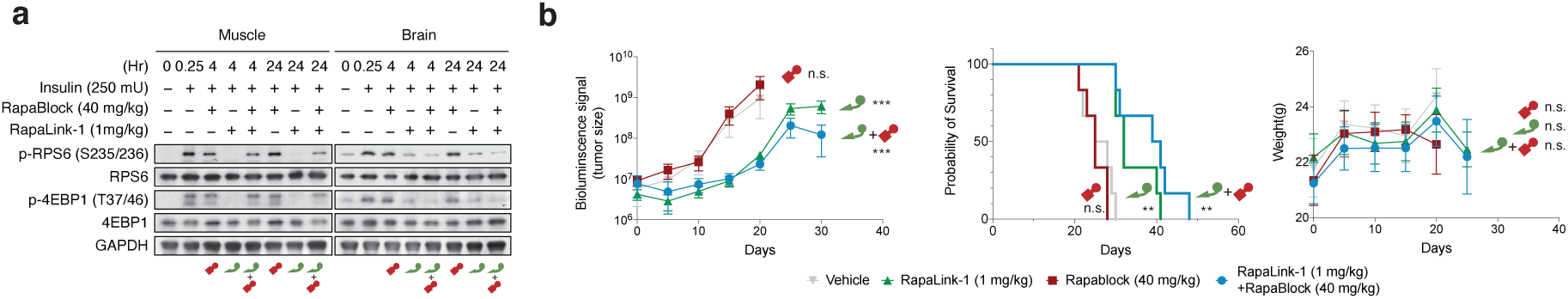
Cotreatment with RapaLink-1 and RapaBlock allows brain-specific inhibition of mTOR. **a**, Mice were treated intraperitoneally with a single dose of RapaLink-1 (1 mg/kg), RapaBlock (40 mg/kg), or a combination of both. mTOR activity was stimulated with insulin (250 mU) 15 minutes before tissue dissection. Whole brain and skeletal muscle were analyzed by immunoblot (*n* = 3 except for the following: no-insulin group, *n* = 1; insulin-only group, *n* = 2. Results are shown for one animal from each group). **b**, Mice bearing luciferase-expressing orthotopic glioblastoma xenografts (U87MG-Luc) were randomized to four different groups (*n* = 7) and treated intraperitoneally every 5 days: (1) vehicle; (2) RapaBlock (40 mg/kg); (3) RapaLink-1 (1 mg/kg); (4) RapaLink-1 (1 mg/kg) and RapaBlock (40 mg/kg). Tumor size was monitored every 5 days using bioluminescence imaging. Statistical tests were performed for each treatment group versus vehicle-treated group: unpaired Student’s *t*-test (tumor size on day 20); log rank test (survival). n.s., not significant; **, p≤0.01; ***, p≤0.001.

RapaLink-1 has been shown to be efficacious in orthotopic mouse models of glioblastoma but inhibition of mTOR in the periphery does not contribute to efficacy. We asked whether our combination regimen, lacking the ability to inhibit mTOR activity in peripheral tissues, could retain the efficacy of RapaLink-1 in treating glioblastoma. We established orthotopic intracranial xenografts of U87MG cells expressing firefly luciferase in nude mice and treated these mice with i.p. injections of RapaLink-1 (1 mg/kg), RapaBlock (40 mg/kg), or a combination of both every 5 days. All treatments were well tolerated, and no significant changes of body weight were observed (Fig. 4b). RapaLink-1, both as a single-agent or in combination with RapaBlock, significantly suppressed tumor growth and improved survival, whereas RapaBlock alone had no significant effect on either.

Having established a binary therapeutic approach to achieve brain-specific inhibition of mTOR, we wondered whether the same strategy could be adapted to other drugs for which brain-restricted pharmacology would be desirable. One critical challenge is that for this approach to be generalizable, the “active” component (e.g. RapaLink-1) must be dependent on the availability FKBP12. Drugs with this property are rare, few if any besides rapamycin and FK506 are known. Our earlier investigation of RapaLink-1 (Fig. 1) and RapaTAMRA (Extended Data Fig. 1) led us to hypothesize that other bifunctional molecules consisting of a FKBP12-binding moiety and a kinase inhibitor moiety could be conditionally active and amenable to modulation with RapaBlock.

We first tested our hypothesis with GNE7915, a potent and specific inhibitor of leucine-rich repeat kinase 2 (LRRK2) a pre-clinical agent under investigation for the treatment of Parkinson’s disease^35–37^. As gain-of-function mutations of LRRK2 are strongly associated with hereditary and sporadic forms of Parkinson’s diseases, LRRK2 kinase inhibition has been pursued as a potential therapeutic strategy^38,39^. However, recent studies demonstrating that systemic LRRK2 inhibition with small molecule inhibitors induced cytoplasmic vacuolation of type II pneumocytes suggests a potential safety liability for these compounds^40,41^. We synthesized FK-GNE7915 by chemically linking the pharmacophores of FK506 and GNE7915 with a piperazine group (Fig. 5a). In *in vitro* LRRK2 kinase assays, FK-GNE7915 was an inferior inhibitor to GNE7915 (Fig. 5c, IC50s of 107 nM and 3.0 nM, respectively), though its activity was potentiated by including 10 µM FKBP12 in the assay (IC_50_: 21 nM). However, in a cellular assay where we quantified phospho-LRRK2 (S935) levels as a marker for LRRK2 inhibition, FK-GNE7915 was more potent than the parent compound GNE7915 by more than 10-fold (Fig 5d, IC50s of 6.7 nM and 81 nM, respectively). The juxtaposition of these two results suggested a role of the FK506 moiety in the enhanced cellular potency of FK-GNE7915. We reasoned that the high-affinity FK506-FKBP12 interaction can promote the intracellular accumulation of FK-GNE7915 similar to the case of RapaLink-1 and confirmed this using a bifunctional fluorescent probe FK-TAMRA (Extended Data Fig. 4). This feature allowed us to program LRRK2 inhibition by controlling the availability of FKBP12: whereas 100 nM FK-GNE7915 treatment reduced phosphor-LRRK2 (S935) level to 15% of basal level, the combination of 100 nM FK-GNE7915 and 1 µM RapaBlock had no effect on LRRK2 activity (Fig. 5e).

**Figure 5.**
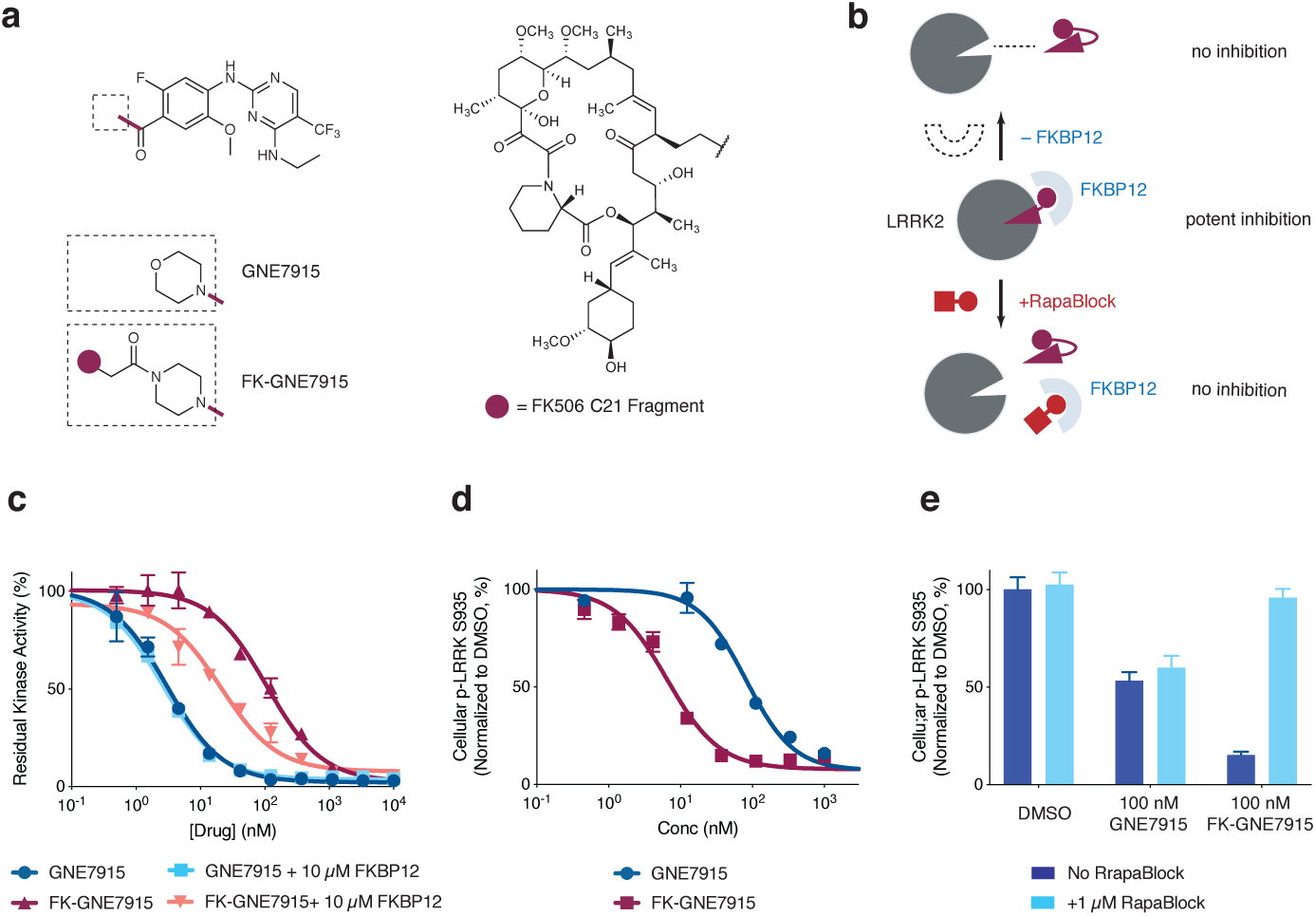
Programmable kinase inhibition with FKBP-dependent kinase inhibitors and RapaBlock. **a**, Structures of GNE7915 and FK-GNE7915. **b**, Proposed working model for FKBP-dependent kinase inhibitors. **c**, kinase inhibition in the absence or presence of supplemented 10 µM recombinant FKBP12 protein. Data is the average of two replicates. **d, e**, RAW264.7 cells were treated with GNE7915, FK-GNE7915, and/or RapaBlock and phospho-LRRK2 (S935) was analyzed by time-resolved FRET using epitopically orthogonal antibodies for LRRK2 and p-LRRK2 (S935). See Supplementary Information for experimental details.

The successful conversion of GNE7915 into a FKBP-dependent LRRK2 inhibitor by simply linking it to an FK506 fragment prompted us to evaluate the generality of this strategy. We explored a number of kinase inhibitors that are being investigated for CNS-diseases: dasatinib (Src family kinase inhibitor, glioblastoma)^42–44^, lapatinib (EGFR/HER2 inhibitor, glioblastoma)^45,46^, and prostetin (MAP4K4 inhibitor, amyotrophic lateral sclerosis and Alzheimer’s disease)^47^. In all three cases, chemically linking the kinase inhibitor to FK506 yielded bifunctional molecules that retained the kinase inhibitory activities of the parent molecules (Extended Data Figs. 5-7). For FK-dasatinib, we also examined its target specificity using both biochemical (Invitrogen SelectScreen Kinase Profiling) and live-cell kinase profiling^48^ and observed an identical spectrum of kinase targets as dasatinib with the exception of DDR2, a known off-target of dasatinib (Extended Data Fig. 5). Despite their large molecular weights, these bifunctional molecules are active in cells with comparable potencies to their parent compounds and sensitive to deactivation by RapaBlock. While these examples represent a limited set of kinase inhibitors with potential CNS disease indications, it is conceivable more drugs can be similarly configured with programmable pharmacology without losing cellular potency.

## Conclusion

Exploiting the unique functional dependence of rapamycin analogs on FKBP12, we have developed an approach to achieve brain-specific mTOR inhibition through the simultaneous administration of a potent mTOR inhibitor (RapaLink-1) and a cell-permeable, BBB-impermeable ligand of FKBP12 (RapaBlock). Tissue-restricted mTOR inhibition enabled by this binary pharmacology strategy reduces the toxicity in peripheral tissues but maintains therapeutic benefits of RapaLink-1 in glioblastoma xenograft models. On the basis of these findings, it seems reasonable to anticipate that the same drug combination may be of broader value in other CNS diseases driven by dysregulated mTOR activity such as Alcohol Use Disorder.

The applicability of our approach extends beyond mTOR inhibition. We show that chemically linking ATP-site kinase inhibitors to FK506 through solvent-exposed groups leads to a new class of cell-permeable kinase inhibitors whose activity depends on the abundant endogenous protein FKBP12. These inhibitors are characterized by their ability to mediate the formation of a ternary complex of the drug, the target kinase and FKBP12, as well as their amenability to activity modulation by RapaBlock. Further *in vivo* studies and medicinal chemistry are necessary to assess and optimize these compounds on a case-by-case basis, but our initial investigations show that it is feasible to attain brain-selective kinase inhibition of LRRK2 using a FKBP-dependent kinase inhibitor (FK-GNE7915) and RapaBlock.

We have investigated the mechanism by which RapaBlock controls the cellular activity of RapaLink-1 as well as other FKBP-dependent kinase inhibitors. Our current data has revealed at least two roles of cellular FKBP12 in the function of RapaLink-1 (and other FKBP-binding compounds). First, we have shown that FKBP12 serves as a reservoir to retain and accumulate RapaLink-1 inside the cell, achieving exceptional cellular concentration. Second, for most FKBP-dependent kinase inhibitors we have investigated, FKBP12 improves their potency in cell-free assays. Though we have not obtained direct evidence, we hypothesize that in aqueous solutions, a bifunctional compound built with flexible linkers between hydrophobic pharmacophores will mainly adopt a binding-incompetent conformation to minimize hydration penalty (“hydrophobic collapse”), and binding of FKBP12 will expose the inhibitor moiety to enable target inhibition. RapaBlock impedes both of these processes by occupying the ligand-binding site of FKBP12. While further investigation is clearly necessary to elucidate how these high molecular weight compounds enter cells and whether FKBP12 participates in the binding interaction with the target protein, we believe this system provides a generalizable approach for programmable kinase inhibition.

Combining two pharmaceutical agents to achieve tissue-selective therapeutic effects has been previously employed in drugs approved for clinical use or in development. Examples include Levodopa/Carbidopa (BBB-permeable dopamine precursor/BBB-impermeable dopa decarboxylase inhibitor) for Parkinson’s Disease^49^, conjugated estrogens/bazedoxifene (BBB-permeable estrogen/BBB-impermeable estrogen receptor modulator) for post-menopausal hot flashes and osteoporosis^50^, and donepezil/solifenacin (BBB-permeable acetylcholinesterase inhibitor/BBB-impermeable anticholinergic) for Alzheimer’s disease^51^. Our approach differs from these precedents in that it does not involve two drugs with counteracting effects on the same target or pathway; instead, RapaBlock controls tissue-specificity by directly attenuating the activity of the kinase inhibitor. The present system therefore has the advantage of being adaptable for a multitude of targets, only requiring an invariant RapaBlock molecule and a FKBP12-dependent inhibitor that can be readily designed based on the structures of FK506 and lead compounds. While we have focused on protein kinases in this study, it seems reasonable to expect that the approach is also applicable to other classes of therapeutic targets, such as GTPases and histone modification enzymes.

## Acknowledgements

We thank Douglas Wassarman for discussions. Z.Z. is a Damon Runyon Fellow supported by the Damon Runyon Cancer Research Foundation (DRG-2281-17). K.M.S, Q.W.F. and W.A.W. acknowledge NIH 1R01CA221969. K.M.S, and W.A.W. acknowledge the Samuel Waxman Cancer Research Foundation. W.A.W. acknowledges NIH U01CA217864 and Cancer Research UK A28592. K.M.S. acknowledges the Michael J. Fox Foundation P0536220, The Mark Foundation for Cancer Research and the Howard Hughes Medical Institute.

## Author Contributions

Z.Z., Q.F., W.A.W and K.M.S. conceived the project, designed and analyzed the experiments, and wrote the manuscript. Z.Z., Q.F., X.L. and K.J.L. performed the laboratory experiments. W.A.W. supervised the *in vivo* experiments.

## Supplementary Information

is available for this paper.

## Author Information

Correspondence and requests for materials should be addressed to K.M.S.

---

**Extended Data Figure 1.**
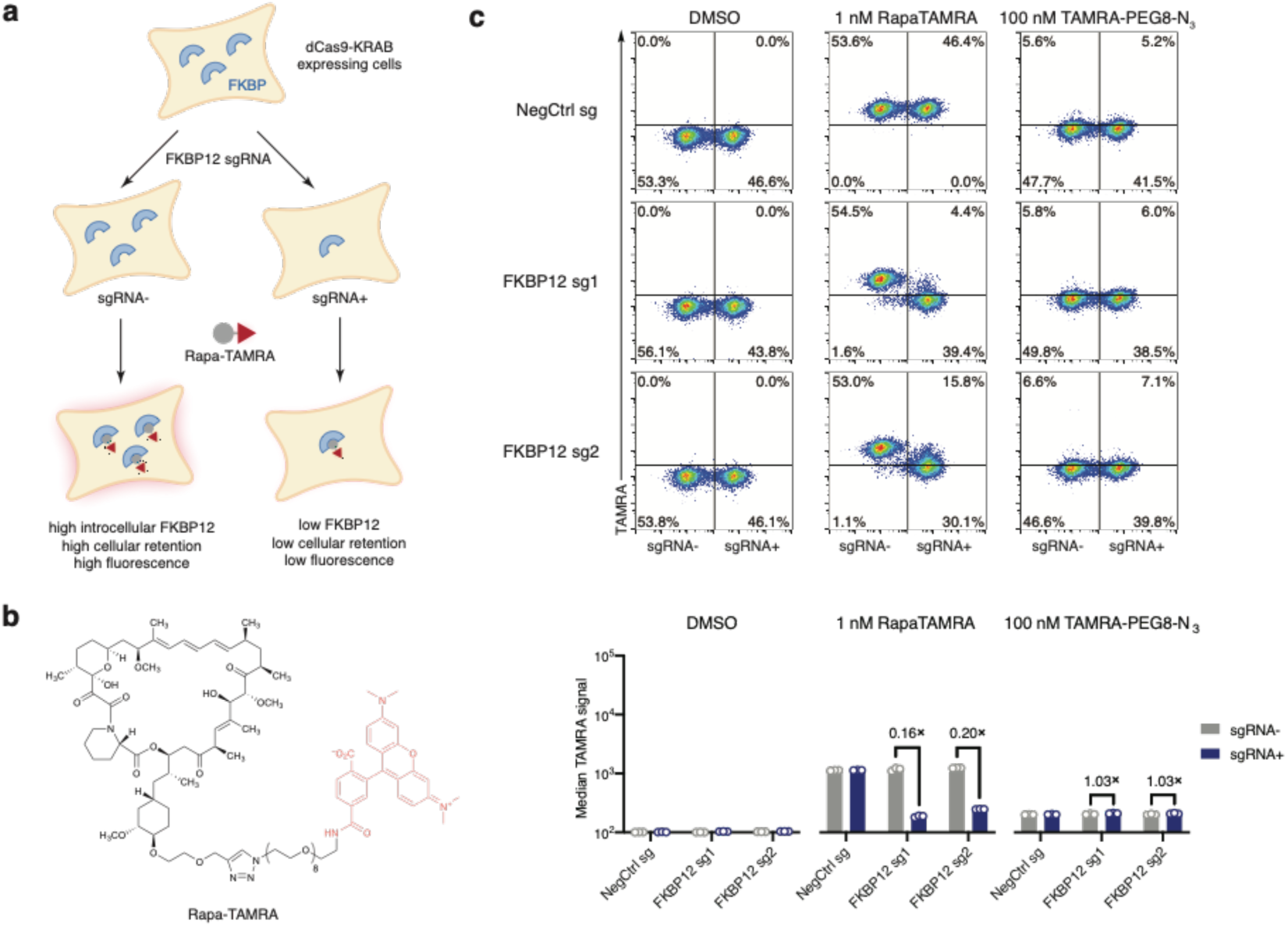
FKBP12-Rapamycin interaction contributes to cellular accumulation of a fluorescent analog of RapaLink-1. **a**, Illustration of the flow cytometry-based assay to assess cellular accumulation of TAMRA compounds. **b**, Structure of a fluorescent probe Rapa-TAMRA. **c**, FKBP12 knockdown decreases cellular retention of Rapa-TAMRA, but not TAMRA-PEG8-N3. See Supplementary Information for gating strategy.

**Extended Data Figure 2.**
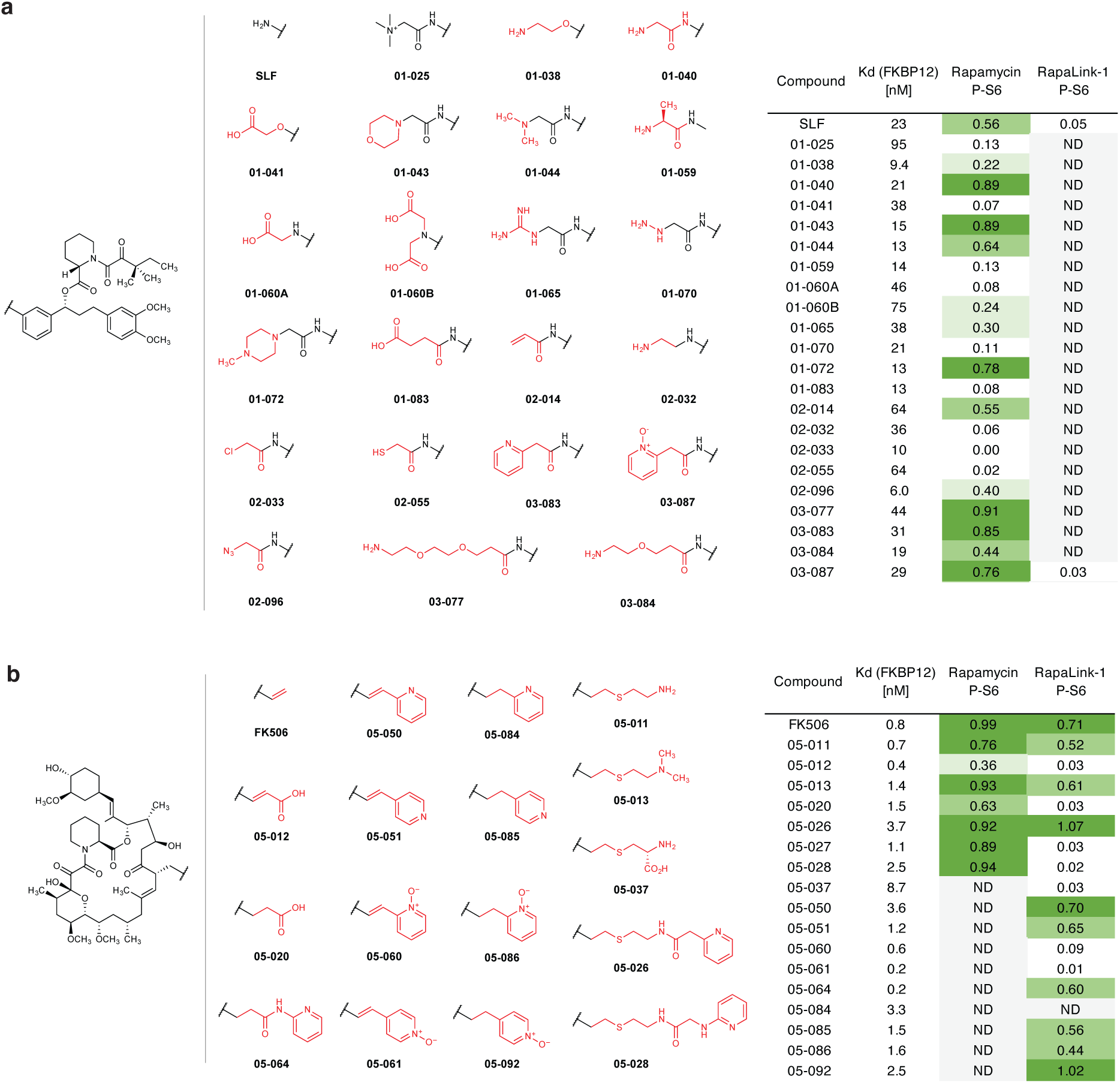
Structures polar FKBP12 ligands synthesized (**a**, SLF derivatives; **b**, FK506 derivatives). Listed in the table are their affinity to recombinant FKBP12 (fluorescence polarization assay) and their efficacy of blocking mTOR inhibition by rapamycin or RapaLink-1 (assessed by western blot analysis of p-S6 level after treatment of MCF7 cells with a combination of 10 nM Rapamycin/Rapalink + 10 µM candidate compound for 24 h. P-S6 level is quantified as fraction of DMSO control). ND, not determined.

**Extended Data Figure 3.**
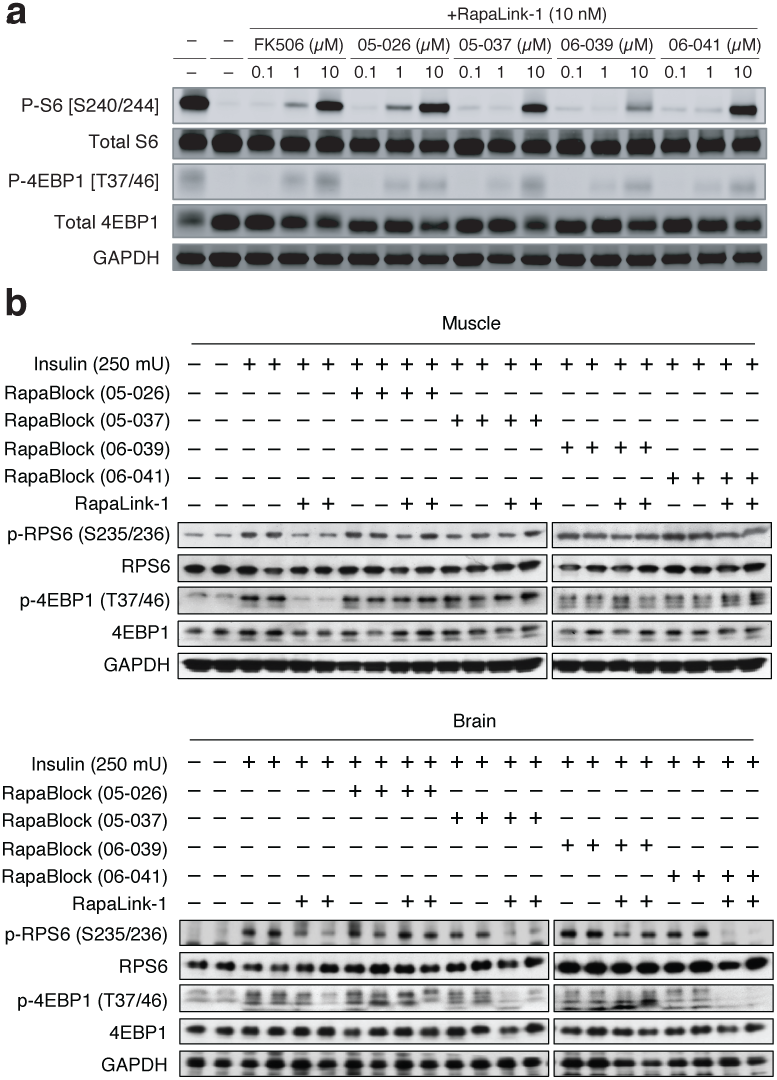
Four candidate RapaBlock molecules were evaluated in cells (a) and *in vivo* (b).

**Extended Data Figure 4.**
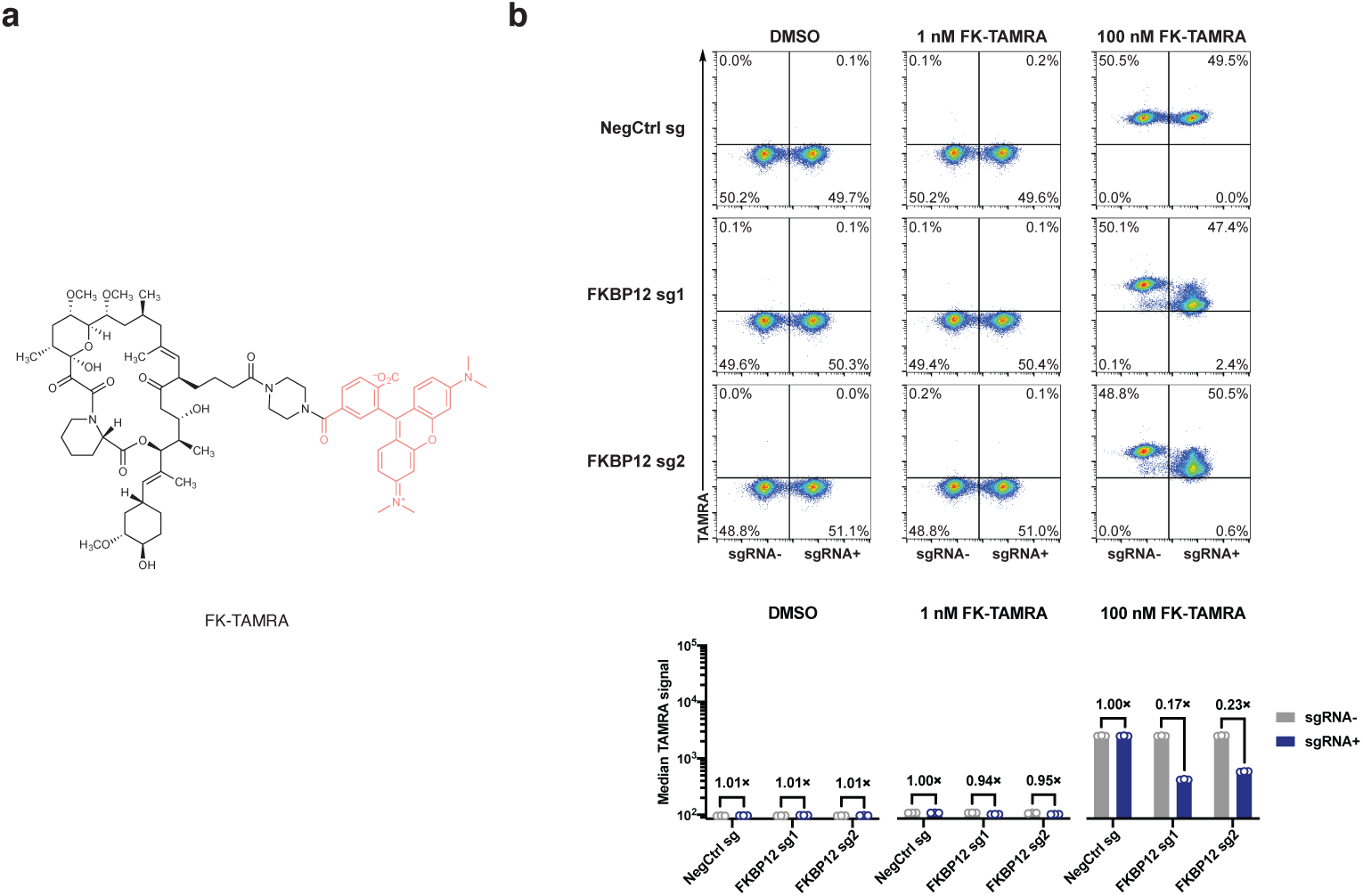
(Related to Extended Data Figure 1) FKBP12-FK506 interaction contributes to cellular accumulation of a fluorescence analog of FK506. **a**, Structure of a fluorescent probe FK-TAMRA. **b**, FKBP12 knockdown decreases cellular retention of FK-TAMRA. See Supplementary Information for gating strategy.

**Extended Data Figure 5.**
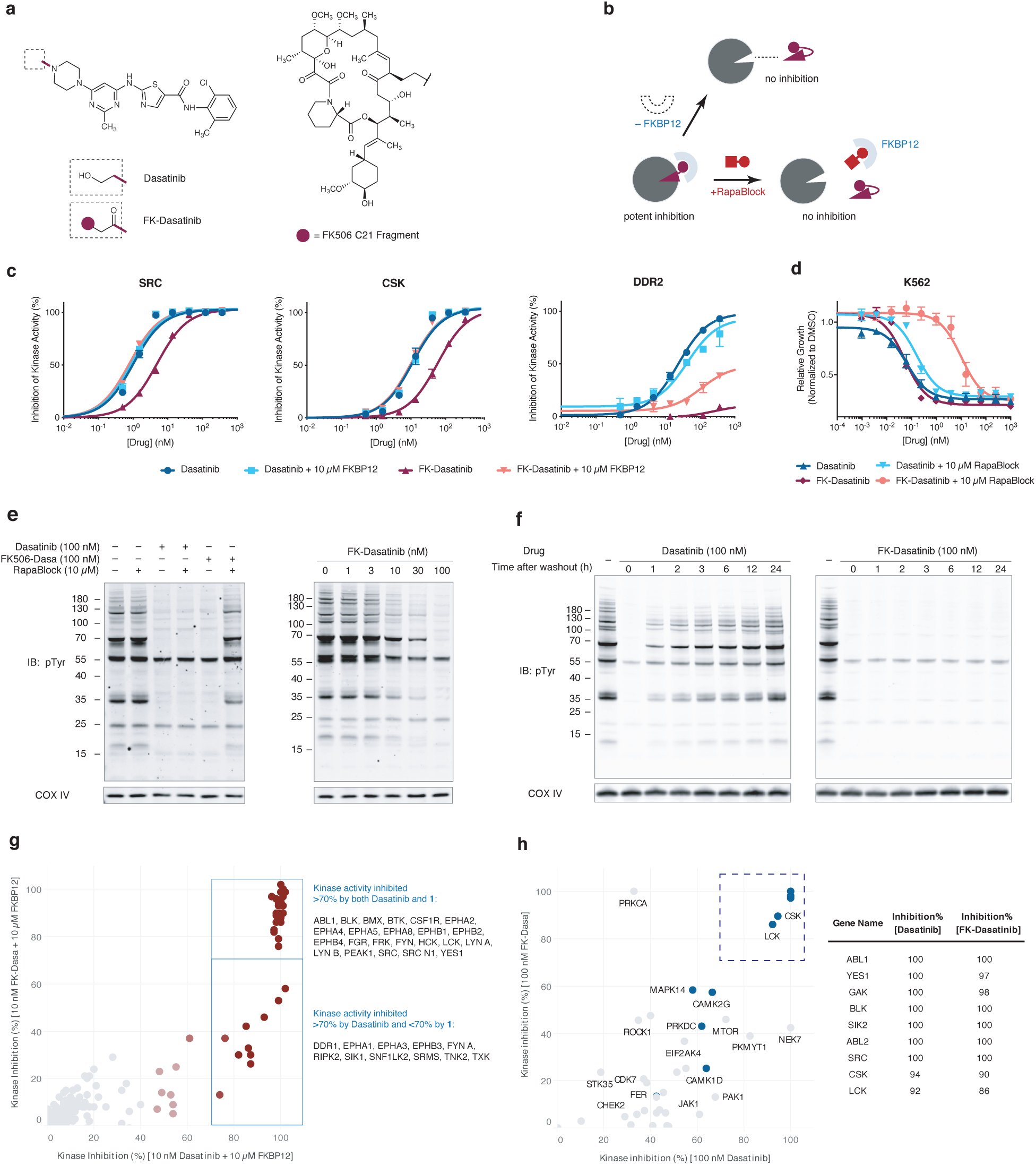
FK-Dasatinib is a FKBP12-dependent Src family kinase inhibitor with long cellular retention time. **a**, Structures of Dasatinib and FK-Dasatinib. **b**, Proposed working model for FKBP-dependent kinase inhibitors. **c**, Inhibition of Src, Csk and DDR2 kinases by Dasatinib and FK-Dasatinib in the absence or presence of supplemented 10 µM recombinant FKBP12 protein. Data is the average of two replicates. **d**, Inhibition of K562 cell proliferation by Dasatinib and FK-Dasatinib, in the presence or absence of 10 µM RapaBlock. Data is the average of three replicates. **e**, Jurkat cells were stimulated with anti-CD3 antibody (OKT3) in the presence of dasatinib, FK-dasatinib, and/or RapaBlock and analyzed by immunoblot. **f**, Jurkat cells were pulse-treated with Dasatinib (100 nM) or FK-Dasatinib (100 nM) for 1 h, then drugs were washed out and cells were incubated in drug-free media for various amounts of time, stimulated with anti-CD3 antibody (OKT3) for 5 min and analyzed by immunoblot. **g**, inhibition of 476 purified kinases by Dasatinib (10 nM) or FK-Dasatinib (10 nM) in the presence of 10 µM recombinant FKBP12 protein. **h**, kinase profiling of Dasatinib (100 nM Dasatinib) or FK-Dasatinib (100 nM) in Jurkat cells using the covalent occupancy probe XO44. See Supplementary Information for experimental details.

**Extended Data Figure 6.**
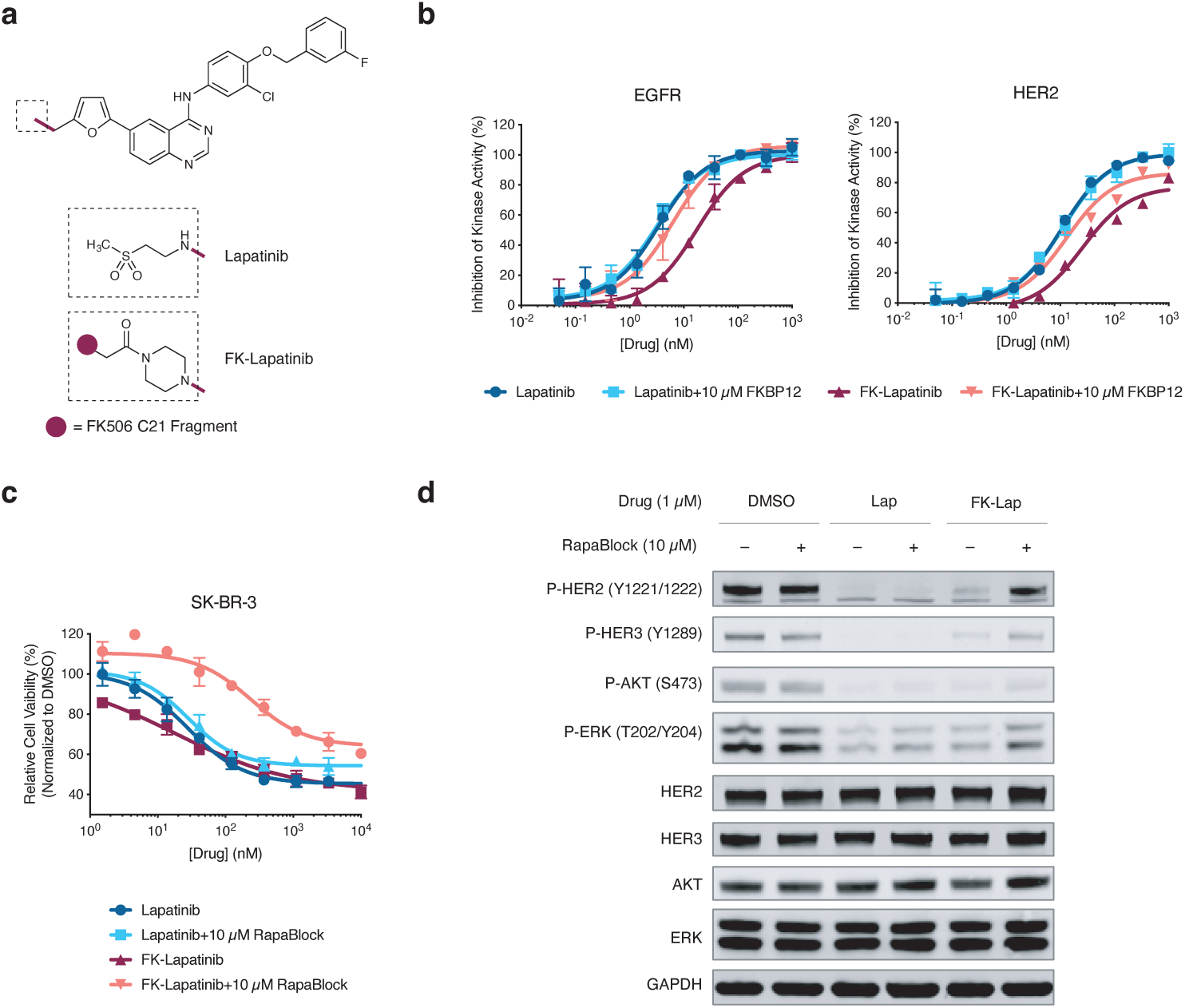
FK-Lapatinib is a FKBP12-dependent EGFR/HER2 kinase inhibitor. **a**, Chemical structures of Lapatinib and FK-Lapatinib. **b**, Inhibition of EGFR and HER2 kinase activity by Lapatinib and FK-Lapatinib in the absence or presence of supplemented 10 µM recombinant FKBP12 protein. Data is the average of two replicates. **c**, Inhibition of proliferation of SK-BR3 cells by lapatinib and FK-lapatinib in the presence of absence of 10 µM RapaBlock. Data is the average of three replicates. **d**, SK-BR3 cells were treated with Lapatinib, FK-Lapatinib, and/or RapaBlock for 1 h and analyzed by immunoblot.

**Extended Data Figure 7.**
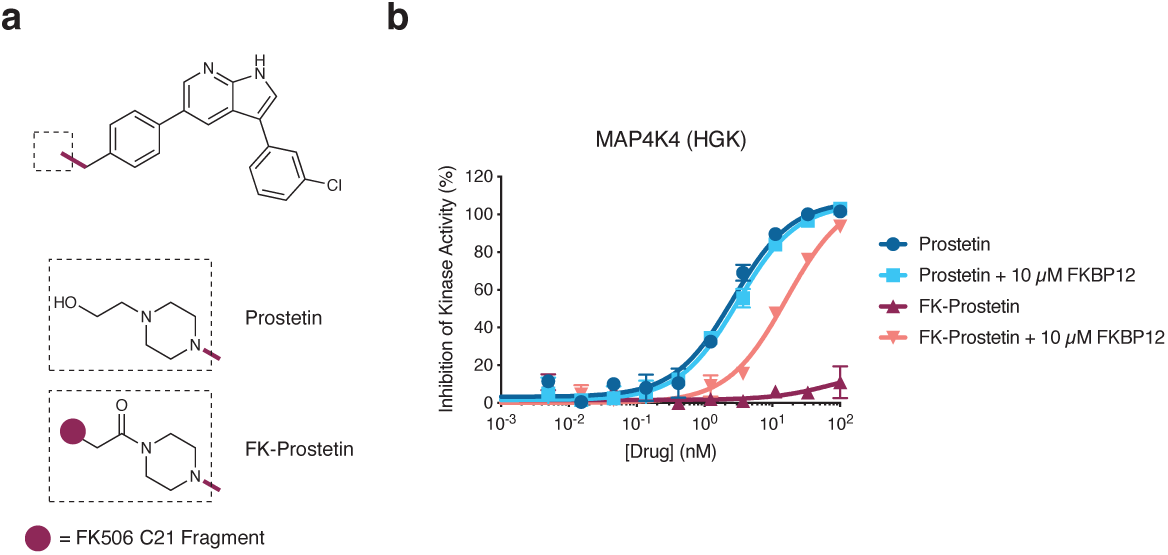
FK-Prostetin is a FKBP12-dependent MAP4K4(HGK) kinase inhibitor. **a**, Chemical structures of Prostetin and FK-Prostetin. **b**, Inhibition of MAP4K4(HGK) kinase activity by Lapatinib and FK-Lapatinib in the absence or presence of supplemented 10 µM recombinant FKBP12 protein. Data is the average of two replicates.

**Extended Data Figure 8.**
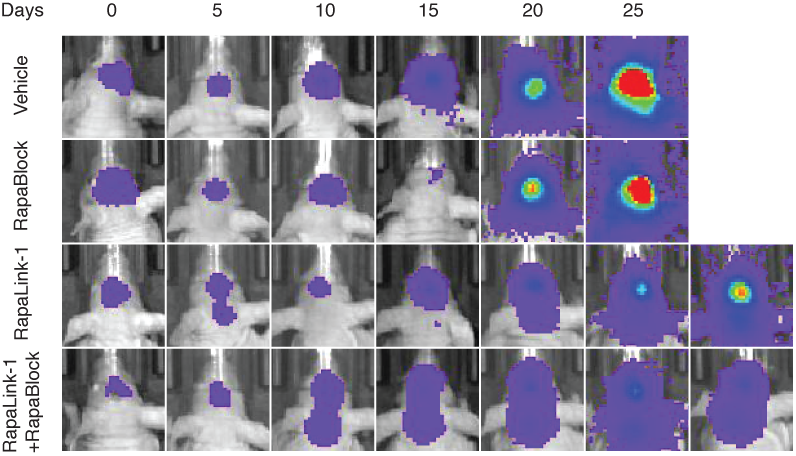
Exemplary bioluminescence images of mice bearing intracranial glioblastoma xenografts. U87MG cells expressing firefly luciferase were injected intracranially into BALB/c^nu/nu^ mice. After tumor establishment, mice were sorted into four groups and treated by i.p. injections every 5 days of vehicle, RapaBlock (1 mg/kg), RapaBlock (40 mg/kg), or a combination of RapaLink-1 and RapaBlock (1 mg/kg and 40 mg/kg, respectively). Bioluminescence imaging of tumor-bearing mice was obtained at days shown (day 0 was start of treatment), using identical imaging conditions.

