## Supporting Information for "Brain-Restricted mTOR Inhibition with Binary Pharmacology"

#### Supplementary Figure(s)

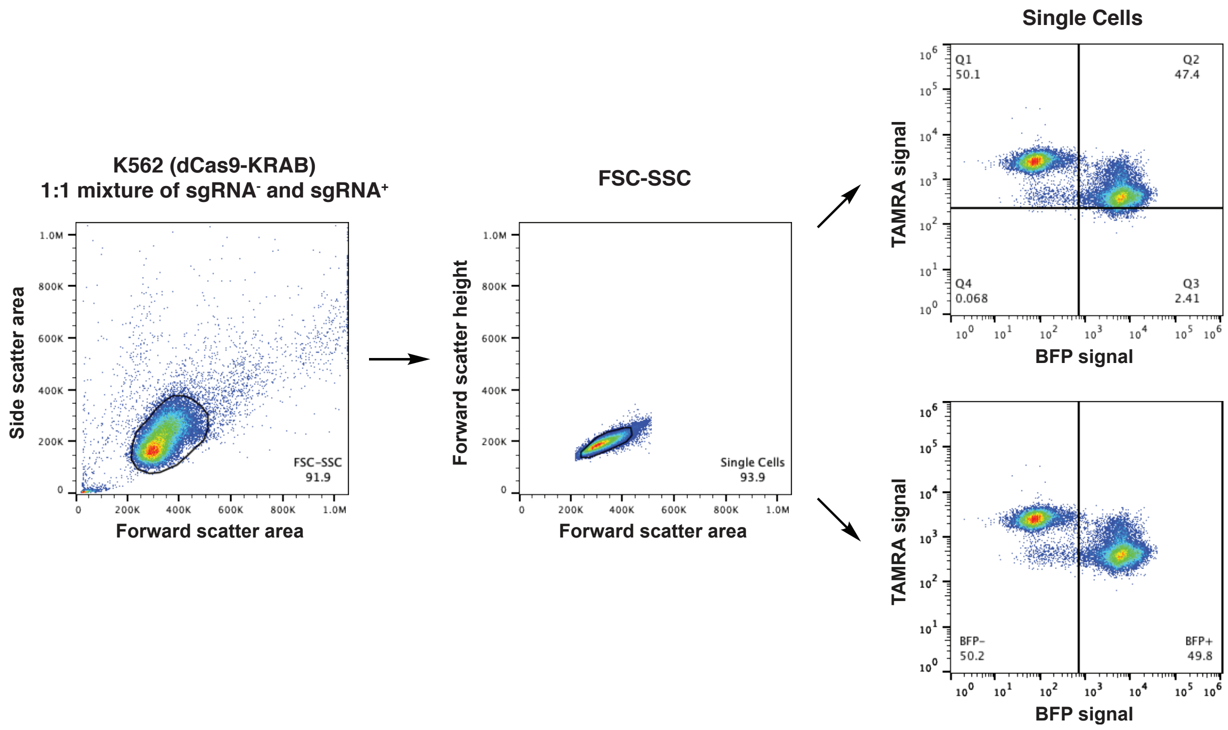

**Figure S1**. Gating strategy for the cellular dye (TAMRA) retention assay.

##### Expression and Purification of Recombinant FKBP12

DNA sequences encoding human FKBP12 were synthesized by Twist Biosciences and cloned into pET47b vector using standard molecular biology techniques (the sequences for these proteins are attached at the end of this document).

Protein expression was performed in BL21(DE3) *E. coli* strain. Briefly, chemically competent BL21(DE3) cells were transformed with pET47b-FKBP12 and grown on LB agar plates containing 50 µg/mL kanamycin at 37 ºC. A single colony was used to inoculate a culture at 37 ºC, 220 rpm in terrific broth containing 50 µg/mL kanamycin. When the optical density reached 0.6, protein expression was induced by the addition of IPTG to 1 mM. After 2 h at 37 ºC, the cells were pelleted by centrifugation (6,500 x g, 10 min) and lysed in lysis buffer [20 mM Tris 8.0, 500 mM NaCl, 5 mM imidazole] with a high-pressure homogenizer (Microfluidics, Westwood, MA). The lysate was clarified by high-speed centrifugation (19,000 x g, 15 min) and the supernatant was used in subsequent purification by immobilized metal affinity chromatography (IMAC). His-tagged FKBP12 was captured by incubation with Co-TALON resin (Clonetech, Takara Bio USA, 4 mL slurry/liter culture) at 4 ºC for 1 h with constant end-to-end mixing. The loaded beads were then washed with lysis buffer (50 mL/liter culture) and the protein was eluted with elution buffer [20 mM Tris 8.0, 500 mM NaCl, 300 mM imidazole]. The His-tag was cleaved with

His-tagged HRV 3C Protease (Clonetech, Takara Bio USA, 5 U/liter culture) at 4 ºC until LC-MS analysis of the reaction mixture indicated >95% cleavage. The reaction mixture was concentrated using an 10K MWCO centrifugal concentrator (Amicon-15, Millipore) to 20 mg/mL and purified by size exclusion chromatography on a Superdex 75 10/300 GL column (GE Healthcare Life Sciences) with SEC Buffer [20 mM HEPES 7.5, 150 mM NaCl]. Fractions containing pure FKBP12 protein were pooled and concentrated to 20 mg/mL and stored at –78 ºC. In our hands, this protocol gives a typical yield of 10-20 mg/liter culture for FKBP12.

##### Determination of Compound Binding Affinity to FKBP12

Compound binding affinity was determined using a competition fluorescence polarization assay. A fluorescent tracer molecule based on rapamycin (FITC-Rapa) was synthesized in house. The assay buffer was 20 mM HEPES 7.5, 0.01% Triton X-100. The Kd of the tracer molecule for FKBP12 was first determined by measuring fluorescence polarization (excitation 485 nm, emission 535 nm) at various protein concentrations and fitting the curve to a quadratic binding model. To measure compound binding affinity, mixtures with the following composition were prepared in duplicate in 96-well black opaque plates (Corning 3915): 0.5 nM FITC-Rapa, 1 nM FKBP12, 5% DMSO, 5 μM–0.08 nM of test compound, 200 μL total volume. Fluorescence polarization was measured on a TECAN Spark 20M plate reader (excitation 485 nm, emission 535 nm). Data were fitted to a three-parameter sigmoidal curve to derive $IC_{50}$ values. Ki of the compounds were calculated using a tool provided by Dr. Shaomeng Wang's lab (http://www.umich.edu/~shaomengwanglab/software/calc_ki/index.html).

##### Cell Culture

MCF7 cells were obtained from ATCC (HTB-22) and maintained in 1:1 DMEM:F12 (Gibco) + 10% heat-inactivated fetal bovine serum (FBS, Axenia Biologix) supplemented with 4 mM L-glutamine, 100 U/mL penicillin and 100 U/mL streptomycin (Gibco). SK-BR-3 cells were obtained from ATCC (HTB-30) and maintained in McCoy's 5A (Gibco) + 10% heat-inactivated FBS supplemented with 2 mM L-glutamine, 100 U/mL penicillin and 100 U/mL streptomycin (Gibco). K562 CRISPRi cells were a gift from Dr. Luke Gilbert and maintained in RPMI 1640 (Gibco) + 10% heat-inactivated FBS supplemented with 2 mM L-glutamine, 100 U/mL penicillin, 100 U/mL streptomycin (Gibco) and 0.1% Pluronic F-68 (Gibco). RAW264.7 cells were obtained from ATCC (TIB-71) and maintained in DMEM (Gibco) + 10% heat-inactivated FBS supplemented with 2 mM L-glutamine, 100 U/mL penicillin and 100 U/mL streptomycin (Gibco). Jurkat cells were obtained from ATCC (TIB-152) and maintained in RPMI 1640 (Gibco) + 10% heat-inactivated FBS supplemented with 2 mM L-glutamine, 100 U/mL penicillin and 100 U/mL streptomycin (Gibco). Jurkat-Lucia NFAT cells were obtained from InvivoGen and maintained in IMDM

(Gibco) + 10% heat-inactivated FBS supplemented with 2 mM L-glutamine, 100 U/mL penicillin, 100 U/mL streptomycin (Gibco) and 100 µg/mL Zeocin (InvivoGen). All cell lines were tested mycoplasma negative using MycoAlert™ Mycoplasma Detection Kit (Lonza). When indicated, cells were treated with drugs at 60-80% confluency at a final DMSO concentration of 1%. At the end of treatment period, cells were placed on ice and washed once with PBS. Unless otherwise indicated, the cells were scraped with a spatula, pelleted by centrifugation (500 x g, 5 min) and lysed in RIPA buffer supplemented with protease and phosphatase inhibitors (cOmplete and phosSTOP, Roche) on ice for 10 min. Lysates were clarified by high-speed centrifugation (19,000 x g, 10 min). Concentrations of lysates were determined with protein BCA assay (Thermo Fisher) and adjusted to 2 mg/mL with additional RIPA buffer. Samples were mixed with 5x SDS Loading Dye and heated at 95 ºC for 5 min.

**Gel Electrophoresis and Western Blot**

Unless otherwise noted, SDS-PAGE was run with Novex 4–12% Bis-Tris gel (Invitrogen) in MES running buffer (Invitrogen) at 200V for 40 min following the manufacturer's instructions. Protein bands were transferred onto 0.45-µm nitrocellulose membranes (Bio-Rad) using a wet-tank transfer apparatus (Bio-Rad Criterion Blotter) in 1x TOWBIN buffer with 10% methanol at 75V for 45 min. Membranes were blocked in 5% BSA–TBST for 1 h at 23 ºC. Primary antibody binding was performed with the indicated antibodies diluted in 5% BSA–TBST at 4 ºC for at least 16 h. After washing the membrane three times with TBST (5 min each wash), secondary antibodies (goat anti-rabbit IgG-IRDye 800 and goat anti-mouse IgG-IRDye 680, Li-COR) were added as solutions in 5% skim milk–TBST at the dilutions recommended by the manufacturer. Secondary antibody binding was allowed to proceed for 1 h at 23 ºC. The membrane was washed three times with TBST (5 min each wash) and imaged on a Li-COR Odyssey fluorescence imager.

**NFAT Activation Assay**

Jurkat-Lucia™ NFAT cells were resuspended to 2 x $10^6$ cells/mL in fresh growth medium and dispensed in 96-well tissue culture plates (Corning 3904, 180 µL/well). DMSO solutions of test compounds at 100x the test concentrations were added and the cells were incubated for 4 h at 37 ºC. A 10x stimulation solution containing phorbol myristate acetate (PMA, 100 ng/mL) and ionomycin (10 µg/mL) were added to each well (20 µL/well) except for the negative control wells, which were supplemented with 20 µL medium. Cells were incubated at 37 ºC for 12 h. 20 µL/well cells were transferred into a white opaque 96-well plate (Corning 3912). TECAN Spark 20M plate reader equipped with an auto-injection system was primed with Lucia™ luciferase substrate solution and set with the following parameters: 50 µL of injection volume, end-point measurement with a 4 second delay time and 0.1 second integration time. Luciferase activity was measured with the settings above and normalized to DMSO-treated, stimulated cells.

**Jurkat Cell Stimulation**

Jurkat cells were cultured and treated as described in Cell Culture section. At the end of treatment period, 1.5 mL aliquot of cells were pelleted (500 x g, 5 min) and washed once with serum-free RPMI (1 mL). The supernatant was removed, and the cells were resuspended in 100 µL 5 µg/mL OKT3 (Invitrogen, functional grade) in RPMI and immediately placed in a 37-ºC water bath. At 5 min, 25 µL 5x SDS Loading Buffer was added and mixed quickly with the cells. The sample was sonicated at 30% power output for 60 s (1-s on, 1-s off) using Qsonica Q500 Sonicator to shear the DNA. The samples were heated at 95 ºC for 5 min and used for SDS-PAGE.

**Phospho-LRRK2 Quantification by TR-FRET**

Phospho-LRRK2 (S935) was quantified in drug-treated cells using Phospho-LRRK2 (Ser935) cellular kit (Cisbio) following the manufacturer's instructions. RAW264.7 (2 x $10^5$ cells/mL) cells were plated in 6-well tissue culture plates (2 mL/well) 24 h prior to treatment. Cells were treated with compounds at the indicated concentrations for 2 h. Medium was removed by aspiration, and the cells were rinsed with ice-old PBS (1 mL). Cells were lysed with 100 µL Cisbio 1X Lysis Buffer at 23 ºC directly in plate for 30 min. The lysates were transferred into microcentrifuge tubes and clarified by centrifugation (19,000 x g, 10 min). 16 µL clarified lysate was dispensed into one well of a low-volume, round-bottom 384-well plate, and 4 µL LRRK2/p-LRRK2 (S935) antibody master mix (Cisbio) was added to each well. The place was incubated at 23 ºC for 4 h. Time-resolved fluorescence was read on a TECAN Spark 20M plate reader with the following parameters:

Lag time: 60 µs
Integration time: 500 µs
Read A: Excitation filter 320(25) nm, Emission filter 610(25) nm, Gain 130
Read B: Excitation filter 320(25) nm, Emission filter 665(8) nm, Gain 165

TR-FRET signal was calculated as the ratio fluorescence intensity [Read B]/[Read A].

**DNA transfections and lentivirus production**

HEK293T cells were transfected with standard packaging vectors using the TransIT-LT1 Transfection Reagent (Mirus Bio). Viral supernatant was collected 2 to 3 days after transfection, filtered through 0.45 µm polyvinylidene difluoride filters (Millipore Sigma), and frozen at -80 °C before transduction.

##### Generation of sgRNA-expressing CRISPRi cell lines

sgRNA protospacers targeting *GAL4-4* (NegCtrl sg GAACGACTAGTTAGGCGTGTA) and *FKBP12* (FKBP12 sg1 GACGGCTCTGCCTAGTACCT and FKBP12 sg2 GCCCAGGAGACGGTGAGTAG) (ref PMC5094855) were cloned into pCRISPRia-v2 (marked with a puromycin resistance cassette and BFP, Addgene #84832). Briefly, complementary synthetic oligonucleotides (Integrated DNA Technologies) with flanking BstXI and BlpI restriction sites were annealed and ligated with BstXI/BlpI-digested pCRISPRia-v2. The sgRNA expression vectors were packaged into lentivirus as described above. Knockdown cells were generated by transducing K562 CRISPRi (sgRNA-) cells with the sgRNA expression vectors at an MOI of <1 (20-40% transduction rate) with 8 µg/mL polybrene. Beginning the second day following lentiviral addition, transduced (sgRNA+) cells were selected using 2 µg/mL puromycin (Gibco) until each cell population stably reached ≥95% BFP+ (405 nm excitation laser, 440/50 nm emission filter) by flow cytometry on an Attune NxT (Thermo Fisher Scientific).

##### Cell viability assay

Cells were seeded into 96-well white flat bottom plates (1,000 cells/well) (Corning) and incubated overnight. Cells were treated with the indicated compounds in a nine-point threefold dilution series (100 µL final volume) and incubated for 72 h. In some conditions, 10 µM FK506 or RapaBlock was uniformly added to the dilution series. Cell viability was assessed using a commercial CellTiter-Glo (CTG) luminescence-based assay (Promega). Briefly, the 96-well plates were equilibrated to room temperature before the addition of diluted CTG reagent (100 µL) (1:4 CTG reagent:PBS). Plates were placed on an orbital shaker for 30 min before recording luminescence using a Spark (Tecan) plate reader.

##### Cellular Dye (TAMRA) Retention Assay

Untransduced (No virus or sgRNA-) and sgRNA expression vector-transduced (sgRNA+) K562 CRISPRi cells were mixed in a 1:1 ratio, plated in 48-well plates, and incubated overnight. Cell mixtures were treated with the indicated TAMRA-linked compounds (300 µL final volume) and incubated for 24 h. Plates were placed over ice and 200 µL of each compound-cell mixture was transferred to a 96-well U-bottom plate. Cells were pelleted at 500*g* for 5 min, and subsequently washed 2× with ice-cold FACS buffer (PBS + 1% BSA + 0.1% $NaN_3$). Cells were resuspended in 200 µL of ice-cold FACS buffer and assessed using an Attune NxT (Thermo Fisher Scientific). Relative uptake of TAMRA-linked compounds was determined by comparing TAMRA fluorescence (561 nm excitation laser, 585/16 emission filter) between sgRNA- and sgRNA+ cells within each well.

#### *In Vitro* Kinase Inhibition Assay

*In Vitro* kinase inhibition assays were performed by SelectScreen Services (Thermo Fisher Scientific) using either Z'-LYTE (mTOR, Src, Csk, EGFR, HER2, HGK), Adapta (LRRK2) or LanthaScreen (DDR2) assay format (standardized protocol is provided here: http://assets.thermofisher.com/TFS-Assets/BID/Methods-&-Protocols/20180123_SSBK_Customer_Protocol_and_Assay_Conditions.pdf). When indicated, recombinant FKBP12 protein was added to the assay mixture to a final concentration of 10 μM.

#### List of Antibodies

| Target | Supplier | Identifier | Dilution |
|---|---|---|---|
| P-AKT [S473] | Cell Signaling Technology | 4060 | 1:1000 |
| AKT | Cell Signaling Technology | 2920 | 1:1000 |
| P-S6 [S240/S244] | Cell Signaling Technology | 5364 | 1:2000 |
| P-S6 [S235/S236] | Cell Signaling Technology | 4858 | 1:2000 |
| S6 | Cell Signaling Technology | 2217 | 1:1000 |
| P-4EBP1 [T37/46] | Cell Signaling Technology | 2855 | 1:1000 |
| 4EBP1 | Cell Signaling Technology | 9644 | 1:1000 |
| FKBP12 | abcam | 58072 | 1:1000 |
| Actin | Proteintech | 60008-1-Ig | 1:50000 |
| GAPDH | Proteintech | 60004-1-Ig | 1:50000 |
| P-Tyr (4G10) | EMD Millipore | 05-321 | 1:1000 |
| COX IV | Cell Signaling Technology | 4850 | 1:1000 |
| P-ERK [T202/Y204] | Cell Signaling Technology | 9101 | 1:1000 |
| Total ERK | Cell Signaling Technology | 4695 | 1:1000 |
| P-HER2 [Y1221/1222] | Cell Signaling Technology | 2243 | 1:1000 |
| P-HER3 [Y1289] | Cell Signaling Technology | 2842 | 1:1000 |
| HER2 | Cell Signaling Technology | 4290 | 1:1000 |
| HER3 | Cell Signaling Technology | 4754 | 1:1000 |

#### List of Buffer Composition

| Name | Composition |
|---|---|
| RIPA Buffer | 25 mM Tris 7.4<br>150 mM NaCl<br>0.1% SDS<br>1% NP-40<br>0.5% sodium deoxycholate |
| FKBP12 FP Assay Buffer | 20 mM HEPES 7.5<br>0.01% Triton X-100 |

#### Chemical Synthesis

General Notes

General Experiment Procedure

All reactions were performed in oven-dried glassware fitted with rubber septa under a positive pressure of argon, unless otherwise noted. Air- and moisture-sensitive liquids were transferred via syringe. Solutions were concentrated by rotary evaporation at or below 40 °C. Analytical thin-layer chromatography (TLC) was performed using glass plates pre-coated with silica gel (0.25-mm, 60-Å pore size, 230−400 mesh, Merck KGA) impregnated with a fluorescent indicator (254 nm). TLC plates were visualized by exposure to ultraviolet light (UV), then were stained by submersion in a 10% solution of phosphomolybdic acid (PMA) in ethanol or an acidic ethanolic solution of *p*-anisaldehyde,[1] followed by brief heating on a hot plate. Flash column chromatography was performed with Teledyne ISCO CombiFlash EZ Prep chromatography system, employing pre-packed silica gel cartridges (Teledyne ISCO RediSep).

Solvents and Reagents

Anhydrous solvents were purchased from Acros Organics. Unless specified below, all chemical reagents were purchased from Sigma-Aldrich and AK Scientific. Commercial solvents and reagents were used as received. FK506 was purchased from LC Laboratories (Woburn, MA).

Instrumentation

Proton nuclear magnetic resonance ($^{1}$H NMR) spectra and carbon nuclear magnetic resonance ($^{13}$C NMR) spectra were recorded on Bruker AvanceIII HD 2-channel instrument (400 MHz/100 MHz) at 23 °C. Proton chemical shifts are expressed in parts per million (ppm, δ scale) and are referenced to residual protium in the NMR solvent ($CHCl_3$: δ 7.26, $D_2HCOD$: δ 3.31). Carbon chemical shifts are expressed in parts per million (ppm, δ scale) and are referenced to the carbon resonance of the NMR solvent ($CDCl_3$: δ 77.0, $CD_3OD$: δ 49.0). Data are represented as follows: chemical shift, multiplicity (s = singlet, d = doublet, t = triplet, q = quartet, dd = doublet of doublets, dt = doublet of triplets, m = multiplet, br = broad, app = apparent), integration, and coupling constant (*J*) in Hertz (Hz). High-resolution mass spectra were obtained using a Waters Xevo G2-XS time-of-flight mass spectrometer. Unless otherwise specified, diastereomeric ratios of products are reported as (major diastereomer) : (sum of minor diastereomers).

Note on rotamers in $^{1}$H and $^{13}$C NMR data:

[1] This solution was prepared by sequential additions of concentrated sulfuric acid (5.0 mL), glacial acetic acid (1.5 mL) and *p*-anisaldehyde (3.7 mL) to absolute ethanol (135 mL) at 23 °C with efficient stirring.

All of the FK506 analogs synthesized here exist as a mixture of two amide rotamers in $CDCl_3$ or $CD_3OD$ (Mierke, D, F.; Schmieder, P.; Karuso, P.; Kessler, H. *Helv. Chim. Acta.* 1991, **74**, 1027–1047.). Due to extensive spectral overlap of the two, the coupling pattern of certain protons can be complicated even if they should display clear splitting patterns in theory. Sometimes, extensive spectral overlap prevents the identification of all peaks of the minor rotamer, and on occasion, of the major rotamer. In this document, only $^1$H NMR peaks of the major rotamer are reported in the best effort of resolving the peaks. $^{13}$C NMR peaks of both rotamers are reported collectively.

Mini-workup

When a mini-workup (A/B) is indicated in the procedure, it was performed as follows: an aliquot (5 μL) of the reaction mixture was retrieved with a glass pipet and added to a plastic vial containing 0.2 mL organic solvent A and 0.2 mL aqueous solution B. The vial was shaken vigorously and allowed to stand until the two layers partitioned. The organic layer was then used for TLC or LC-MS analysis as specified in the procedure.

Monitoring Reaction Progress by LC-MS

When LC-MS analysis of the reaction mixture is indicated in the procedure, it was performed as follows. An aliquot (1 μL) of the reaction mixture (or the organic phase of a mini-workup mixture) was diluted with 100 μL 1:1 acetonitrile:water. 1 μL of the diluted solution was injected onto a Waters Acquity UPLC BEH C18 1.7 μm column and eluted with a linear gradient of 5–95% acetonitrile/water (+0.1% formic acid) over 3.0 min. Chromatograms were recorded with a UV detector set at 254 nm and a time-of-flight mass spectrometer (Waters Xevo G2-XS).

Synthesis of RapaBlock

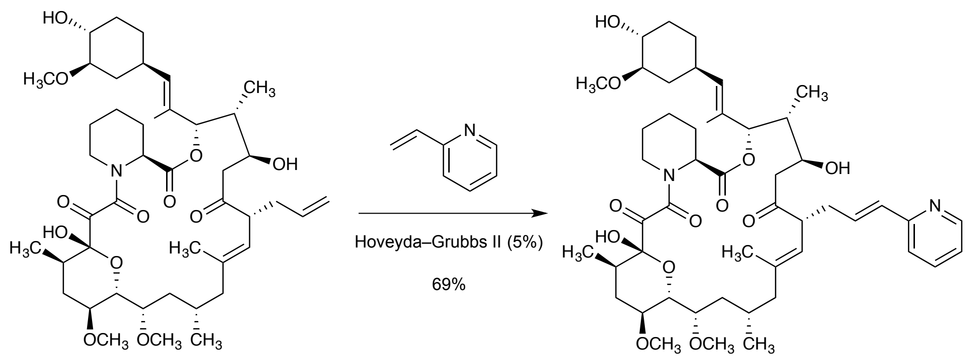

FK506 S1

A 15-ml microwave vial was dried with gentle flame under vacuum. The vial was cooled to 23 °C, flushed with argon, then charged with FK506 (50 mg, 0.062 mmol), DCE (0.62 mL) and a magnetic stir bar. Argon was bubbled through the resulting solution via a 19-gauge needle for 1 min. 2-Vinylpyridine (6.7 µL, 0.062 mmol) was added via pipette, and Grubbs-Hoveyda $2^{nd}$ Gen Catalyst (4.0 mg, 0.0062 mmol) was added in one portion as a solid. The mixture was stirred briefly (giving a bright green solution) before being loaded on a CEM DiscoverSP microwave reactor. Microwave reaction was performed at 100 °C for 30 min with 1 min pre-equilibration. After cooling to 23 °C, the reaction mixture was analyzed by TLC (100% ethyl acetate), which showed formation of an UV-active, more polar spot. The reaction mixture was directly loaded onto a 4-g RediSep (Teledyne ISCO) column. Elution with 100% ethyl acetate gave the product as a yellow solid (38 mg, 69%). 3:2 mixture of rotamers.

$^{1}$H NMR (400 MHz, Chloroform-*d*) δ 8.55 – 8.47 (m, 1H), 7.66 – 7.56 (m, 1H), 7.24 (tt, *J* = 8.1, 1.1 Hz, 1H), 7.16 – 7.06 (m, 1H), 6.69 – 6.56 (m, 1H), 6.57 – 6.46 (m, 1H), 5.38 – 5.30 (m, 1H), 5.16 – 5.00 (m, 2H), 4.63 (d, *J* = 5.3 Hz, 1H), 4.44 (d, *J* = 14.1 Hz, 1H), 4.02 – 3.84 (m, 2H), 3.77 – 3.67 (m, 1H), 3.66 – 3.50 (m, 2H), 3.42 (s, 3H), 3.40 (s, 3H), 3.38 – 3.35 (m, 1H), 3.31 (s, 3H), 3.07 – 2.96 (m, 3H), 2.86 – 2.64 (m, 3H), 2.51 – 2.23 (m, 4H), 2.24 – 1.85 (m, 8H), 1.71 – 1.60 (m, 6H), 1.60 – 1.31 (m, 8H), 1.14 – 1.03 (m, 2H), 1.01 (d, *J* = 6.3 Hz, 3H), 0.94 (d, *J* = 6.7 Hz, 3H), 0.89 (d, *J* = 7.2 Hz, 3H).

$^{13}$C NMR (100 MHz, Chloroform-*d*) δ 212.69, 212.32, 196.06, 192.70, 168.90, 168.70, 165.78, 164.63, 155.65, 155.48, 149.32, 149.27, 140.05, 139.21, 136.43, 132.48, 132.34, 131.89, 131.84, 131.77, 131.66, 129.63, 129.55, 122.41, 122.26, 121.82, 121.76, 121.07, 121.04, 98.57, 97.04, 84.12, 77.83, 77.20, 76.47, 75.07, 73.59, 73.50, 73.47, 72.68, 72.16, 70.10, 68.96, 65.82, 57.51, 56.93, 56.61, 56.58, 56.54, 56.29, 56.09, 52.73, 52.67, 52.61, 48.52, 48.31, 44.09, 43.84, 43.24, 40.47, 39.98, 39.22, 35.32, 34.86, 34.82, 34.70, 34.57, 34.40, 34.12, 33.56, 32.76, 32.67, 32.52, 31.17, 30.58, 27.62, 26.17, 25.98, 24.50, 21.12, 20.83, 20.40, 19.37, 16.21, 15.97, 15.95, 15.82, 15.24, 14.22, 14.16, 9.75, 9.48.

HRMS (ESI): Calcd for $(C_{49}H_{72}N_2O_{12} + H)^+$: 881.5163, Found: 881.5207

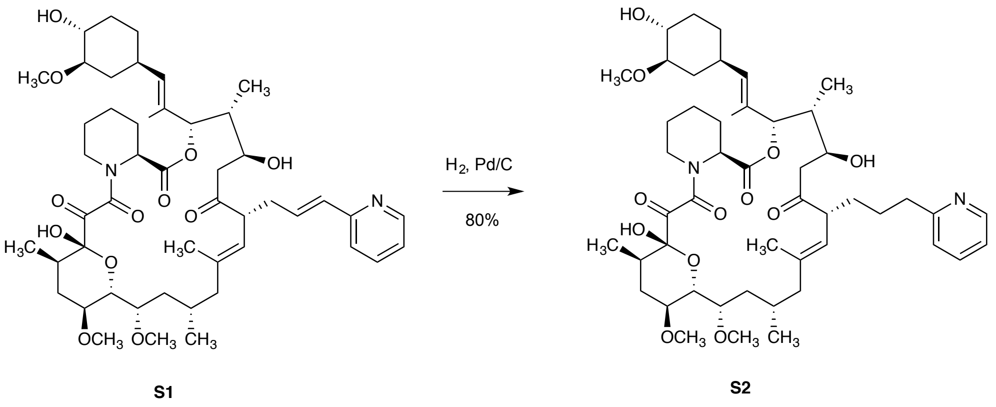

A 20-mL vial was charged with **S1** (20 mg, 0.020 mmol), Ethyl acetate (1.0 mL) and Palladium on carbon (10 wt%, 2.4 mg). The vial was briefly purged with argon, and then fitted with a rubber septum. Hydrogen was bubbled through the solution via a 19-gauge needle for 5 min, then the mixture was stirred under hydrogen atmosphere at 23 ºC. In a total of 3 h, LC-MS showed full conversion to the desired m/z. The reaction mixture was filtered through a pad of Celite, and the filter cake was rinsed with ethyl acetate (5 mL). The combined filtrate was concentrated to afford the product as a pale-yellow foam (20 mg, 99%). 3:2 mixture of rotamers.

$^1$H NMR (400 MHz, Chloroform-d) δ 8.56 – 8.43 (m, 2H), 7.61 (tt, J = 7.6, 2.3 Hz, 2H), 7.21 – 7.07 (m, 4H), 5.37 (s, 1H), 5.28 – 5.15 (m, 1H), 5.15 – 4.96 (m, 2H), 4.62 (d, J = 5.0 Hz, 1H), 4.45 (d, J = 13.5 Hz, 1H), 4.00 – 3.86 (m, 1H), 3.74 (d, J = 9.8 Hz, 1H), 3.67 – 3.55 (m, 1H), 3.52 – 3.25 (m, 3H), 3.43 (s, 3H), 3.41 (s, 3H), 3.32 (s, 3H), 3.09 – 2.95 (m, 1H), 2.92 – 2.75 (m, 2H), 2.73 – 2.64 (m, 1H), 2.43 – 2.11 (m, 3H), 2.11 – 1.84 (m, 4H), 1.84 – 1.70 (m, 4H), 1.70 – 1.57 (m, 6H), 1.57 – 1.33 (m, 8H), 1.15 – 1.05 (m, 2H), 1.03 (d, J = 6.5 Hz, 3H), 0.96 (d, J = 6.1 Hz, 3H), 0.87 (d, J = 7.2 Hz, 3H).

$^{13}$C NMR (100 MHz, Chloroform-d) δ 213.96, 213.18, 196.35, 192.17, 168.95, 168.22, 166.01, 164.96, 161.79, 161.65, 149.16, 148.94, 139.64, 138.28, 136.48, 136.36, 132.46, 131.80, 129.57, 129.26, 123.24, 122.84, 122.71, 121.04, 98.59, 97.33, 84.13, 77.80, 77.20, 76.68, 76.50, 75.15, 73.68, 73.51, 72.64, 72.18, 70.25, 68.85, 57.54, 56.88, 56.66, 56.59, 56.53, 56.30, 56.06, 52.98, 52.66, 52.09, 48.23, 42.86, 40.41, 39.77, 39.14, 37.97, 37.53, 34.84, 34.72, 33.60, 32.84, 32.64, 31.18, 30.75, 30.61, 29.91, 27.44, 27.33, 27.20, 26.10, 25.94, 24.47, 21.18, 20.83, 20.55, 19.45, 16.35, 16.16, 15.97, 15.65, 14.27, 14.21, 9.74, 9.49.

HRMS (ESI): Calcd for $(C_{49}H_{74}N_2O_{12} + H)^+$: 883.5320, Found: 883.5332

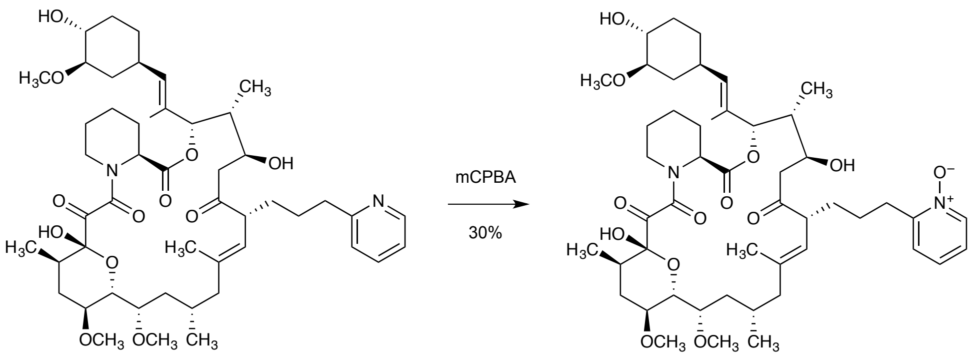

**S2** **RapaBlock**

*m*-CPBA (6.6 mg, 0.025 mmol) was added as a 10% solution (w/v) in dichloromethane (66 µL) to a solution of **S2** (22 mg, 0.025 mmol) in dichloromethane at 0 ºC. The reaction progress was monitored by LC-MS. In a total of 6 h, LC-MS showed full conversion to the desired m/z. The reaction mixture was directly concentrated under reduced pressure. The residue was diluted with 50% acetonitrile–water to a volume of 3.0 mL, and the solution was filtered through a 0.45 µM PTFE syringe filter. The filtrate was purified by reverse-phase HPLC (Waters XBridge C18 column 5 µm particle size 30 x 250 mm, 50–95% acetonitrile–water + 0.1% formic acid, 40 min, 20 mL/min) to afford the product as a white solid (6.6 mg, 30%). 3:2 mixture of rotamers.

$^{1}$H NMR (400 MHz, Chloroform-*d*) δ 8.32 – 8.22 (m, 2H), 8.03 (s, 1H), 7.24 – 7.11 (m, 3H), 5.34 (s, 1H), 5.13 (d, *J* = 9.6 Hz, 1H), 5.10 – 5.04 (m, 1H), 4.78 – 4.67 (m, 1H), 4.58 (d, *J* = 3.3 Hz, 1H), 4.43 (d, *J* = 13.8 Hz, 1H), 3.93 (t, *J* = 10.2 Hz, 1H), 3.77 – 3.68 (m, 1H), 3.64 – 3.52 (m, 1H), 3.41 (s, 3H), 3.39 (s, 3H), 3.48 – 3.23 (m, 3H), 3.30 (s, 3H), 3.06 – 2.91 (m, 3H), 2.83 – 2.73 (m, 1H), 2.73 – 2.65 (m, 1H), 2.41 – 2.23 (m, 3H), 2.22 – 2.08 (m, 3H), 2.06 – 1.94 (m, 2H), 1.94 – 1.71 (m, 6H), 1.71 – 1.50 (m, 6H), 1.50 – 1.30 (m, 8H), 1.06 (d, *J* = 13.4 Hz, 2H), 1.00 (d, *J* = 6.5 Hz, 3H), 0.93 (d, *J* = 5.8 Hz, 3H), 0.86 (d, *J* = 7.2 Hz, 3H).

$^{13}$C NMR (100 MHz, Chloroform-*d*) δ 213.54, 212.84, 196.34, 192.92, 168.99, 168.74, 165.82, 164.97, 152.29, 152.11, 139.85, 139.75, 139.70, 138.52, 132.56, 131.94, 129.57, 129.40, 126.31, 125.97, 125.66, 125.54, 123.55, 123.03, 122.82, 98.59, 97.27, 84.16, 77.76, 77.27, 76.47, 74.82, 73.70, 73.61, 73.54, 73.50, 72.69, 72.21, 70.30, 69.24, 65.85, 57.53, 56.93, 56.69, 56.63, 56.60, 56.57, 56.33, 56.14, 52.77, 52.68, 52.34, 48.37, 44.01, 43.88, 43.21, 40.56, 39.98, 39.17, 35.31, 34.91, 34.87, 34.77, 34.72, 33.60, 32.87, 32.61, 31.26, 30.65, 30.58, 30.32, 27.48, 26.24, 26.15, 26.04, 24.51, 23.77, 23.65, 21.20, 20.93, 20.53, 19.46, 16.26, 16.23, 16.00, 15.85, 15.28, 14.29, 14.27, 9.84, 9.60.

HRMS (ESI): Calcd for $(C_{49}H_{74}N_2O_{13} + H - H_2O)^+$: 881.5164, Found: 881.5150

### $^{1}H$ NMR and $^{13}C$ NMR Spectra

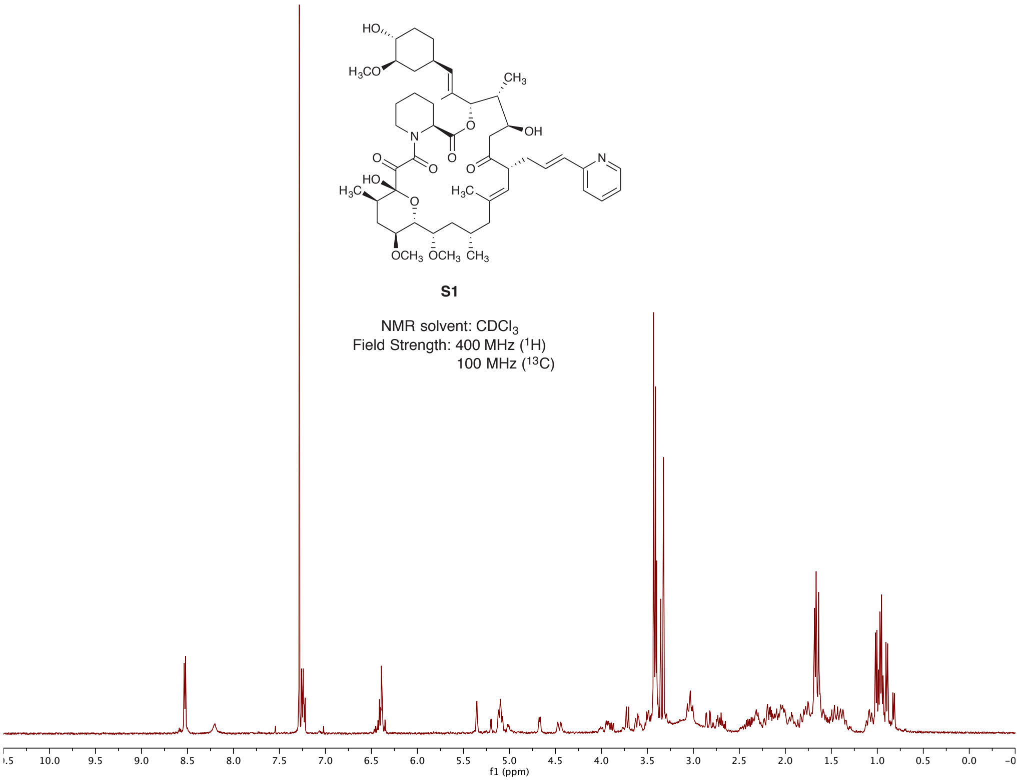

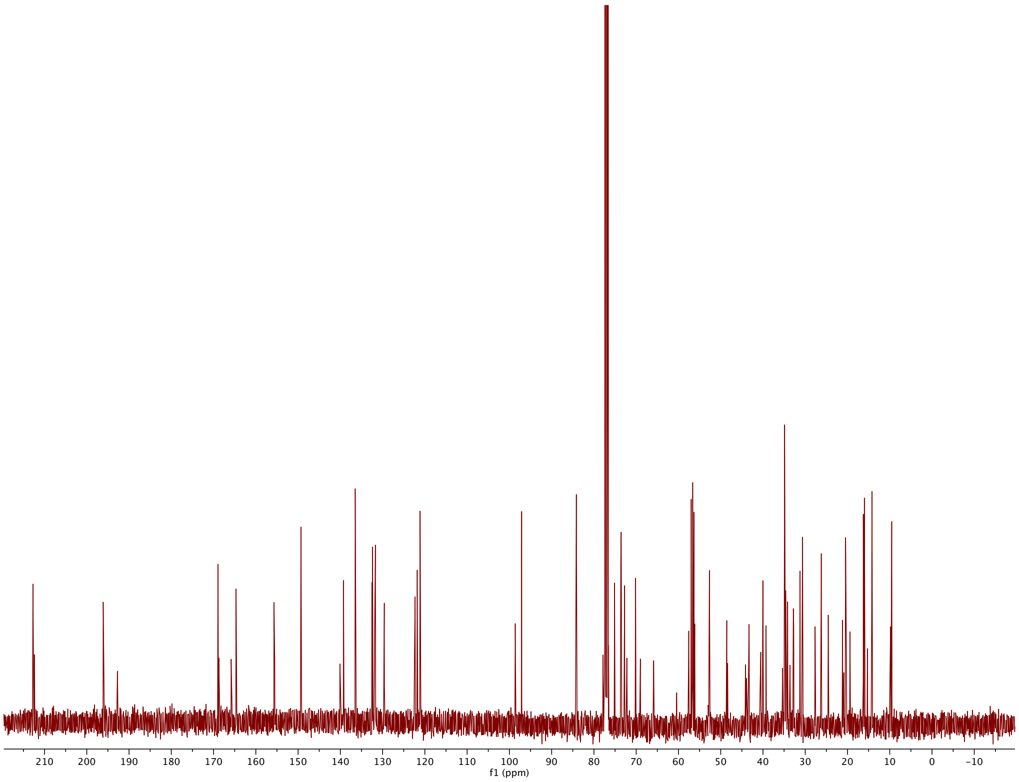

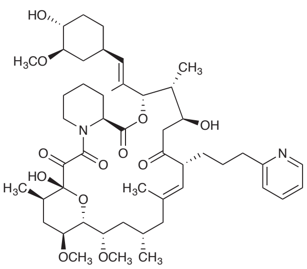

**S2**

NMR solvent: $CDCl_3$
Field Strength: 400 MHz ($^1H$)
100 MHz ($^{13}C$)

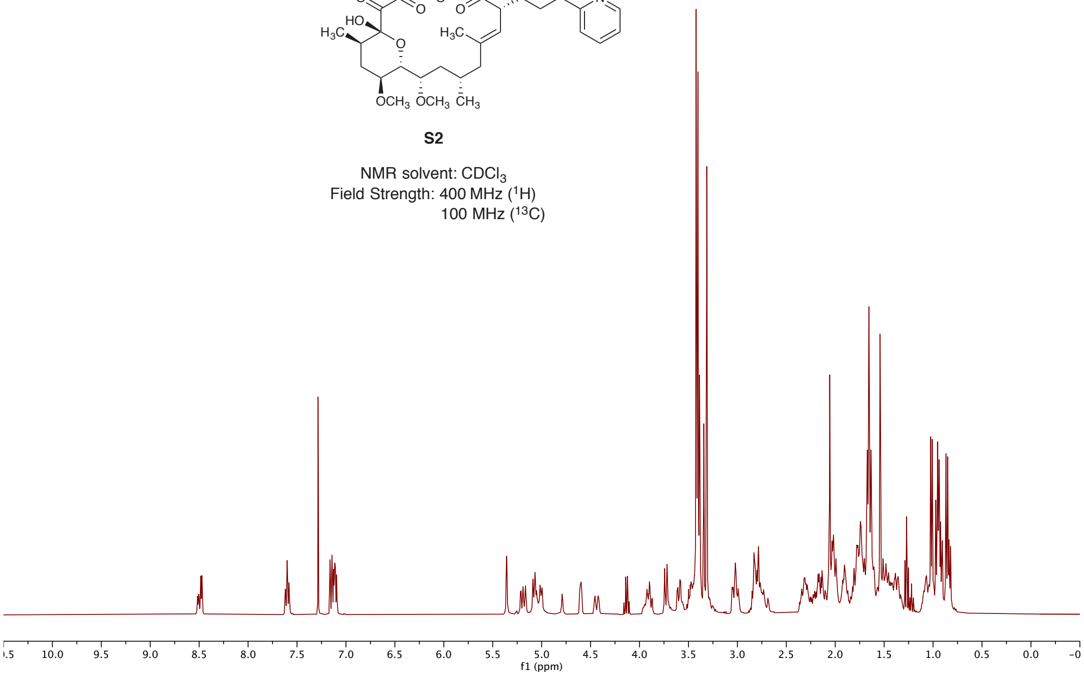

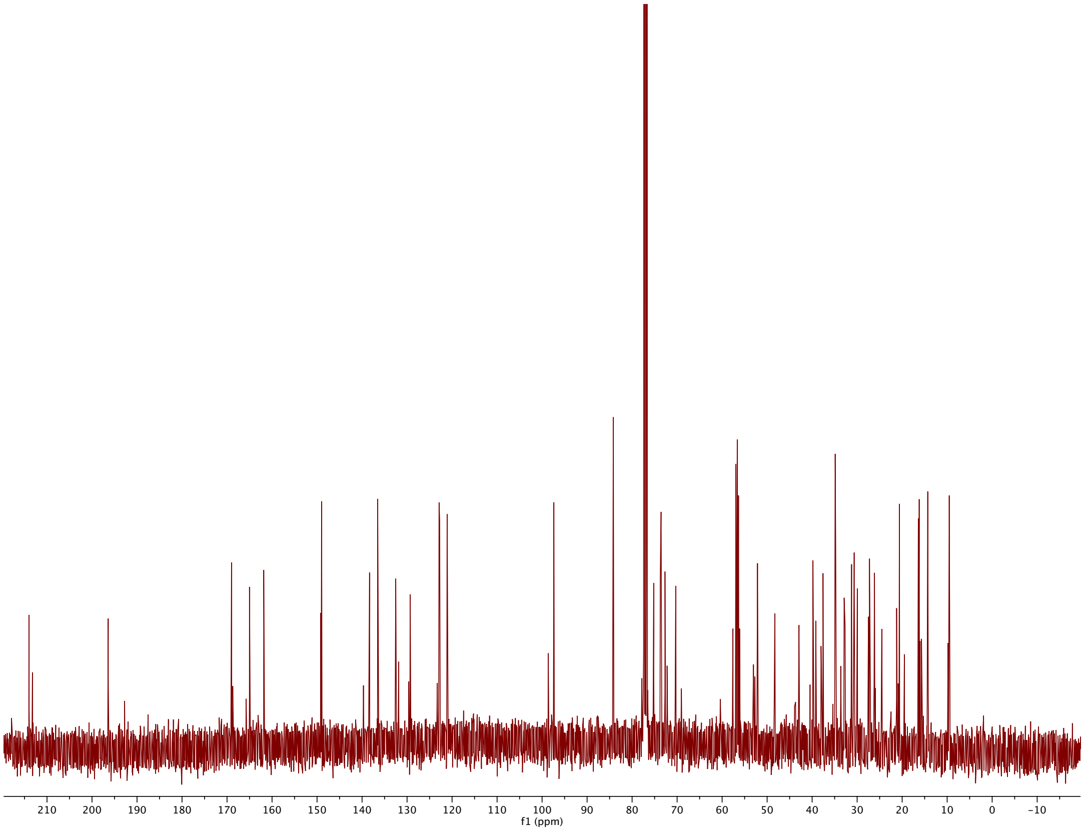

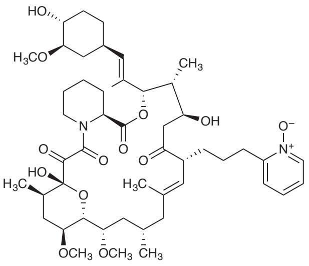

**RapaBlock**

NMR solvent: $CDCl_3$
Field Strength: 400 MHz ($^1H$)
100 MHz ($^{13}C$)

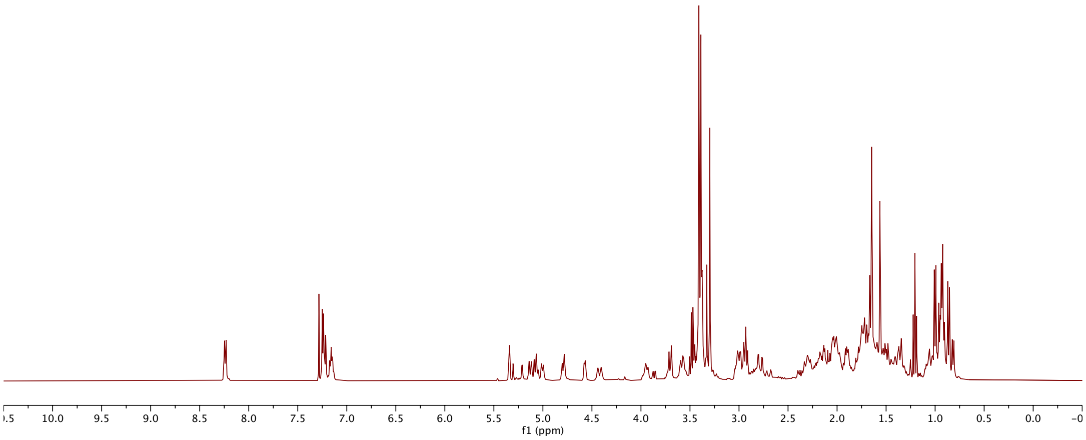

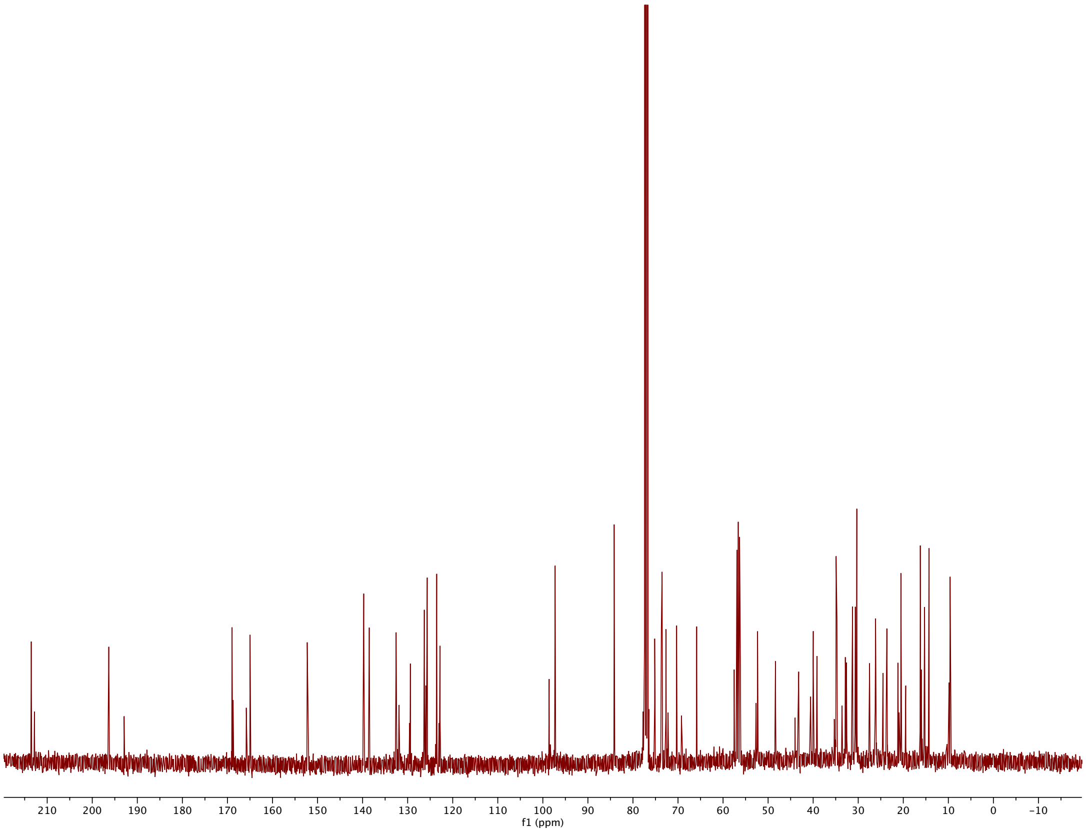
